# Functional plasticity and recurrent cell states of malignant B cells in follicular lymphoma

**DOI:** 10.1101/2022.04.06.487285

**Authors:** Noudjoud Attaf, Chuang Dong, Laurine Gil, Inãki Cervera-Marzal, Tarek Gharsalli, Jean-Marc Navarro, Diana-Laure Mboumba, Lionel Chasson, François Lemonnier, Philippe Gaulard, Sandrine Roulland, Lionel Spinelli, Bertrand Nadel, Pierre Milpied

## Abstract

Follicular lymphoma (FL) derives from malignant transformation of germinal center (GC) B cells. FL malignant B cells are heterogeneous and diverge from their GC B cell-of-origin, but the diversity, function, and location of malignant B cell states remain to be addressed. Based on integrative single-cell RNA-seq, we identified and studied recurrent FL malignant B cell states and dynamics. Most FL B cells spanned a continuum of states from proliferating GC-like to quiescent memory (Mem)-like cell states. That GC-to-Mem axis was the main source of intra-tumor transcriptional heterogeneity. While FL B cell states were independent from subclonal B cell receptor genetics divergence, T follicular helper (T_FH_) cell-derived signals controlled the transition from Mem-like to GC-like states. GC-like, T_FH_-activated and Mem-like FL B cells tended to occupy distinct niches within and around tumor follicles. Our study characterizes novel malignant cell states recurrent in B cell lymphomas, and highlights the functional plasticity of malignant B cells.

## INTRODUCTION

Germinal centers (GCs) are microanatomical structures that form within secondary lymphoid organs upon T-cell dependent B cell activation^1^. Within GCs, antigen-specific B cells undergo clonal expansion, antibody affinity maturation and terminal differentiation into antibody-secreting plasma cells (PC) or recirculating memory (Mem) B cells. CD4^+^ T follicular helper (T_FH_) cells control GC reactions at multiple stages, from GC seeding by activated B cells to affinity-based selection and differentiation into PC and Mem B cells^2^. Upon antigen re-exposure, most Mem B cells differentiate into PC, and some Mem B cells reenter GCs, leading to further diversification of their B cell receptor (BCR) repertoire^3–7^. GCs are dynamic structures where proliferation, somatic mutations and cell death are tightly regulated to promote immunity while avoiding B cell lymphomagenesis^8–10^. Yet, many of such control points may fail, and accordingly, most B cell lymphomas arise from GC-experienced B cells^11–14^.

Follicular lymphoma (FL) is the second most frequent lymphoma in adults and is considered a prototypical GC-derived malignancy. Within malignant lymph nodes, FL B cells are clustered in follicular structures and express typical GC B cell markers CD20, CD10, BCL6, as well as the activation-induced cytidine deaminase AID which is driving Ig somatic hypermutation^15,16^. FL often presents as a disseminated disease at diagnosis and its progression is indolent with a median overall survival of more than 15 years with current therapies^17–19^. Although treatment based on CD20-targeting antibodies and chemotherapy is effective, FL patients often suffer from multiple relapses. Almost 20% of patients progress or relapse within 2 years after treatment initiation, which is associated with poor clinical outcome^20^. Given the heterogeneity of clinical evolution patterns, understanding the biological mechanisms promoting FL dissemination, progression and relapse is a major challenge.

Genetically, FL B cells are characterized by the t(14;18)(q32;q21) chromosomal translocation in approximately 90% of cases^21^, leading to the ectopic expression of the anti-apoptotic BCL-2 protein normally silenced in GC B cells. Other recurrent genetic lesions target the epigenetic modifiers KMT2D (∼80% of cases)^22^ and CREBBP (∼65% of cases)^23^. Studies in human cohorts^24,25^ and mouse models^26^ have shown that FL lymphomagenesis involves a long asymptomatic phase where pre-malignant t(14;18)^+^ B cells get drawn into successive GC reactions, presumably leading to additional genetic hits and post-translational modifications like BCR N-glycosylation^27,28^ which trigger final transformation into the overt FL stage. FL B cells are admixed to stromal and immune cell subsets, and the tumor microenvironment (TME) plays a key role in FL pathogenesis and clinical outcome^29–31^. Among TME components, FL T_FH_ cells sustain the survival and proliferation of FL B cells through the expression of CD40L, IL4, IL21, among other signals^29,31,32^.

FL B cells include quiescent centrocytes resembling light zone GC B cells, and large proliferating centroblasts resembling dark zone GC B cells^33^. Centrocytes usually outnumber centroblasts, but a high density of centroblasts within FL follicles is a marker of more aggressive disease^11^. Bulk transcriptional profiling studies have shown that FL B cells harbor a gene expression profile closely related to light zone GC B cells^34,35^. However, our recent study has revealed that FL B cells are not locked in a specific GC B cell state^36^. In that study, we showed that FL B cell transcriptional states did not follow GC-specific synchronized gene expression patterns. Other single-cell RNA-seq (scRNA-seq) studies have identified that FL B cells express a patient-specific gene expression program, with some degree of intra-patient transcriptional heterogeneity^37,38^. Although genetic heterogeneity may imprint transcriptional and functional heterogeneity^37,38^, our previous study suggested that some FL gene expression states are independent from subclonal genetic divergence and may be recurrent across patients^36^. Those recurrent B cell states may sustain core functions in B cell lymphomas and remain to be characterized.

Here, we have studied intra-tumor heterogeneity and characterized recurrent malignant B cell states in human FL samples using integrative single-cell RNA-seq. Our study identifies novel FL B cell subsets, and reports that intra-tumoral heterogeneity of malignant B cells results mainly from functional plasticity in response to TME cues.

## RESULTS

### Inter-patient heterogeneity in FL B cells

We first analyzed FL B cells and non-malignant B cells with FB5P-seq, a plate-based scRNA-seq method enabling parallel analysis of transcriptome and BCR repertoire in FACS-sorted B cells^39^ (**Fig.1A-B**). Our cell sorting strategies purified PC, Mem and GC B cells subsets from spleen and tonsils, and isotype-restricted CD10^+^ malignant B cells from FL lymph nodes (**Fig.S1A-B)**. We visualized single-cell gene expression data with uniform manifold approximation and projection (UMAP), a dimensionality reduction method that maps closely related cells into clusters^40^. FL B cells clustered on the basis of sample origin, separately from non-malignant GC, Mem and PC cells, and expressed sample-specific monoclonal BCR rearrangements (**Fig.S2**). Only paired diagnosis and relapse samples from FL338 donor mapped closely (**Fig.1C-D**). Non-malignant B cells clustered depending on their GC, Mem or PC type and expressed polyclonal BCR (**Fig.1C-D, S2**). We also performed high-throughput droplet-based 10x 3’ (**Fig.1E-G**) and 10x 5’ scRNA-seq (**Fig.1H-J**) on all live cells from FL cell suspensions. We first analyzed the 10x 3’ and 10x 5’ datasets separately to avoid technology-related batch effects. We defined TME non-B cells, malignant and non-malignant B cells and visualized them in UMAP. Non-malignant B cells and TME cells did not cluster on the basis of donor origin (**Fig.1F-G, I-J**). Conversely, malignant FL B cells displayed strong inter-patient heterogeneity and only paired diagnosis/relapse samples (FL068 and FL338) mapped together (**Fig.1I-J**).

**Figure 1.**
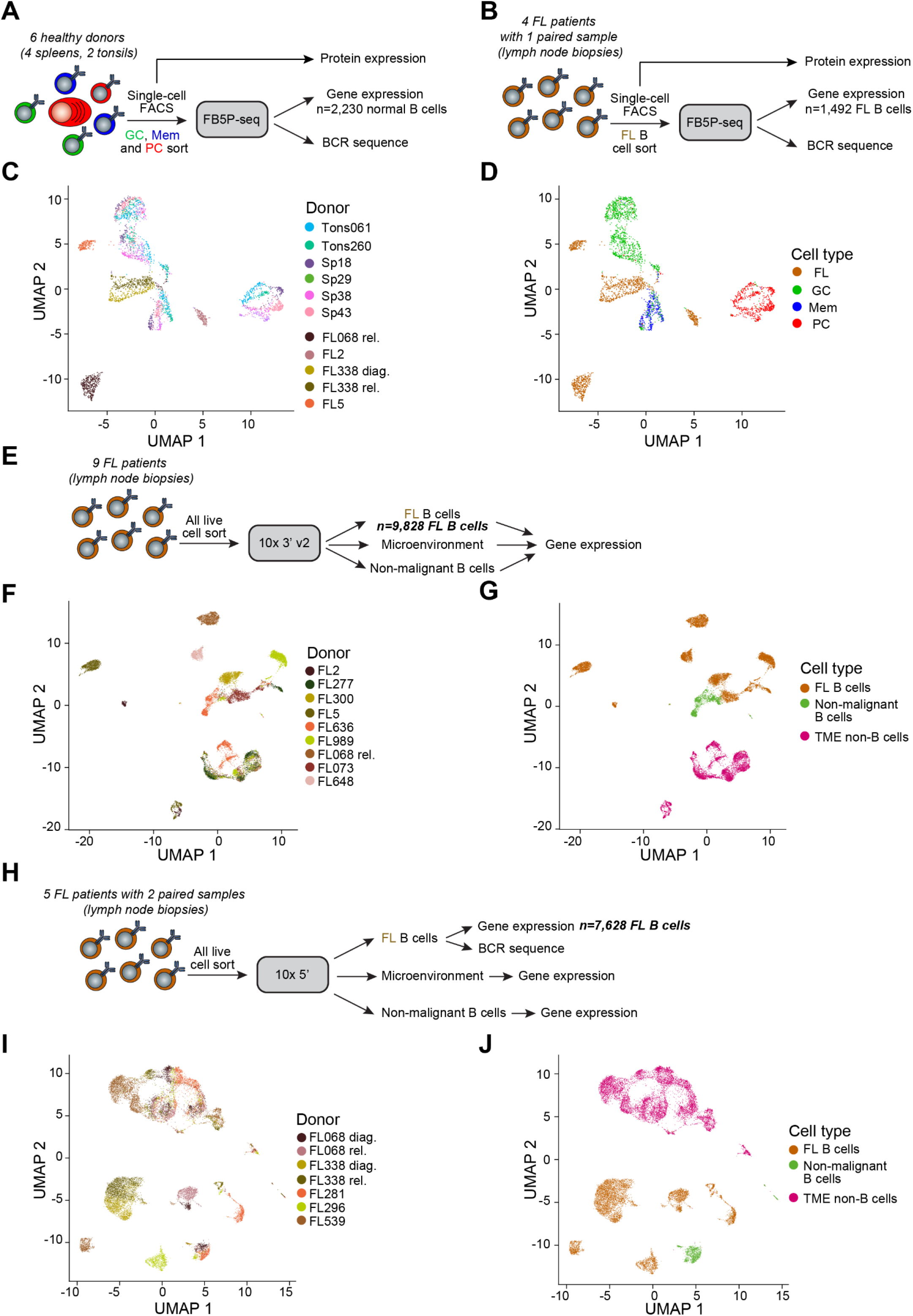
Single-cell RNA-seq identifies inter-patient heterogeneity in FL B cells. **(A)** FB5P-seq experimental workflow on non-malignant human B cells. **(B)** FB5P-seq experimental workflow on human malignant FL B cells. **(C, D)** UMAP embedding of normal and FL B cells analyzed by FB5P-seq. Cells are colored based on donor origin **(C)** or cell type **(D)**. *n*=3,722 B cells. **(E)** High-throughput 3’-end 10x Genomics experimental workflow on total human FL cell suspensions. **(F, G)** UMAP embedding of total FL cell suspensions analyzed by 3’-end 10x Genomics. Cells are colored based on donor origin **(F)** or cell type **(G)**. *n*=15,797 cells. **(H)** High-throughput 5’-end 10x Genomics experimental workflow on total human FL cell suspensions. **(I, J)** UMAP embedding of total FL cell suspensions analyzed by 5’-end 10x Genomics. Cells are colored based on donor origin **(I)** or cell type **(J)**. *n*=18,493 cells.

Thus, malignant FL B cells exhibit patient-specific gene expression profiles distinct from non-malignant B cells.

### Non-malignant B cell signatures

Beyond inter-patient divergence, we sought to identify patterns of intra-patient gene expression heterogeneity in malignant B cells. To characterize potential similarities between malignant and non-malignant B cells, we first computed four lists of differentially expressed genes (DEGs) associated with PC, non-PC, GC and Mem gene expression patterns (detailed in **Computational Methods** and **Fig.S3A-F, Supplementary Data Table 1**). Then for each cell in a scRNA-seq dataset, we reported the expression of genes from a given signature with a single value defined as the signature score. That scoring strategy accurately classified normal GC, Mem and PC subsets analyzed by FB5P-seq (**Fig.S3G-H**) or by 10x 3’ and 10x 5’ scRNA-seq (**Fig.S3I-J**). We then used those signatures to detect malignant B cells expressing gene expression programs close to those of non-malignant B cell types.

### PC-like FL B cells

FL B cells are not expected to resemble antibody-producing PC in most cases, but some cases of FL with plasmacytic differentiation have been described^41^. We first intended to rule out plasmacytic differentiation as a source of transcriptional heterogeneity in malignant B cells. We computed PC and non-PC signature scores for malignant FL B cells in the FB5P-seq, 10x 3’ and 10x 5’ datasets. Almost all FL B cells were transcriptionally distinct from PC cells (**Fig.2A-B**). However, we detected rare FL B cells carrying a PC-like gene expression profile in all datasets. PC-like cell frequency reached 2-3% of malignant FL B cells in 2/15 patients (FL2 and FL539), but those cells were absent or present in frequencies <0.3% in other samples (**Fig.2C, Table S7**). PC-like cells had low expression of *MS4A1* and high expression of PC markers *CD27, XBP1, MZB1* and *PRDM1* (**Fig.2D**). Gene ontology (GO) analysis of genes upregulated in FL539 PC-like cells (**Supplementary Data Table 2**) showed representation of pathways active in PC as a consequence of massive antibody protein synthesis (*e*.*g*. response to endoplasmic reticulum (ER) stress, unfolded protein response, protein N-linked glycosylation) (**Fig.2E**)^42^. BCR-seq analysis showed that PC-like cells expressed the patient-specific malignant monoclonal BCR rearrangement, confirming that they were not non-malignant contaminants. Thus, malignant FL B cells occasionally differentiate into antibody producing PC-like cells, but the vast majority of malignant B cells are not involved in such differentiation.

**Figure 2.**
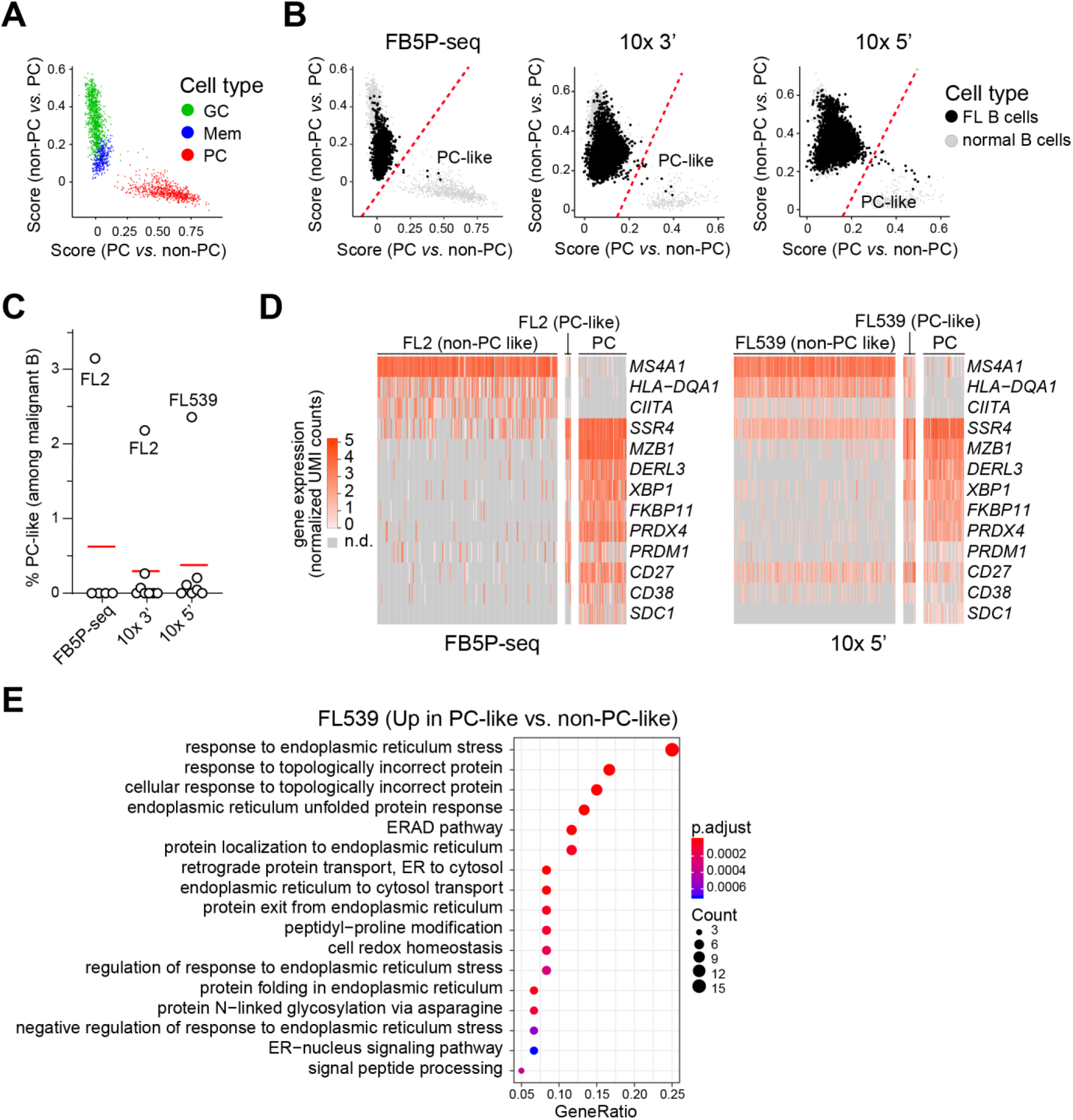
PC-like FL B cells. **(A)** Non-PC *vs*. PC signature score plotted against PC *vs*. non-PC signature score for single normal spleen and tonsil B cells (FB5P-seq) colored by cell type, presented as a scatter plot. **(B)** Non-PC *vs*. PC signature score plotted against PC *vs*. non-PC signature score for malignant FL B cells (black dots), presented as a scatter plot. Grey dots correspond to single normal spleen and tonsil B cells. Red dashed line represents the cut-off used for assigning PC-like cells. From left to right, data from FB5P-seq, 10x 3’ and 10x 5’. **(C)** Per-sample quantification of PC-like FL B cells for each scRNA-seq dataset, calculated as the percentage of PC-like FL B cells among malignant FL B cells in each sample. FL2 and FL539 samples are highlighted. *n*= 5 samples for FB5P-seq. *n*= 8 samples for 10x 3’. *n*= 7 samples for 10x 5’. **(D)** Gene expression heatmap for non-PC like and PC-like FL B cells from FL2 sample (left, FB5P-seq), or from FL539 sample (right, 10x 5’) compared to normal PC. Genes were selected on the basis of their known downregulation (*MS4A1, HLA-DQA1, CIITA*) or upregulation (all other genes) in normal PC cells. **(E)** Gene Ontology analysis of genes upregulated in PC-like FL B cells from FL539 sample. The dot plot shows the number of genes associated with each pathway (dot size, or count) and the adjusted p-values for those terms (color).

### Continuum of states between GC-like and Mem-like cell states

We next focused on the expression of GC and Mem signatures. Only a minority of FL B cells expressed a GC-like profile, some harbored a Mem-like profile, and all FL B cells showed a continuum rather than a clear-cut separation (**Fig.3A-B**). We assigned all FL B cells to one of the GC-like, Mid or Mem-like states based on signature scores thresholds (**Fig.3B**). GC-like and Mem-like cells were detected in all patients with variable proportions among malignant B cells; in most samples the majority of FL B cells were in Mid state (**Fig.3C, Table S7**), expressing low levels of both GC and Mem scores. To account for the continuum of cell states, we also defined a GC-like to Mem-like continuous score (Score_GCtoMem_) that increased from 0 to 1 as FL B cells shifted from GC-like to Mem-like. As expected from the signature scores, GC-like and Mem-like cells expressed GC-specific and Mem-specific marker genes, respectively (**Fig.3D-E**). Proliferating FL B cells (S and G2/M phases, *MKI67* expression) were in the GC-like state (**Fig.3D**). Among recurrent intra-sample DEG markers of FL B cell subsets (**Supplementary Data Table 3**), GC-like FL B cells notably expressed *AICDA, RGS13, CD38, CD81, NANS* and *BCL6*. Mem-like FL B cells were quiescent and expressed extra-follicular homing markers such as *CCR7, CD44, CD69, CXCR4, GPR183, KLF2* and *SELL* genes (**Fig.3D-E**). Consistent with a continuum of states, GC-specific and Mem-specific gene markers exhibited smooth and gradual transitions in expression levels from one state to the other (**Fig.3D-E**).

**Figure 3.**
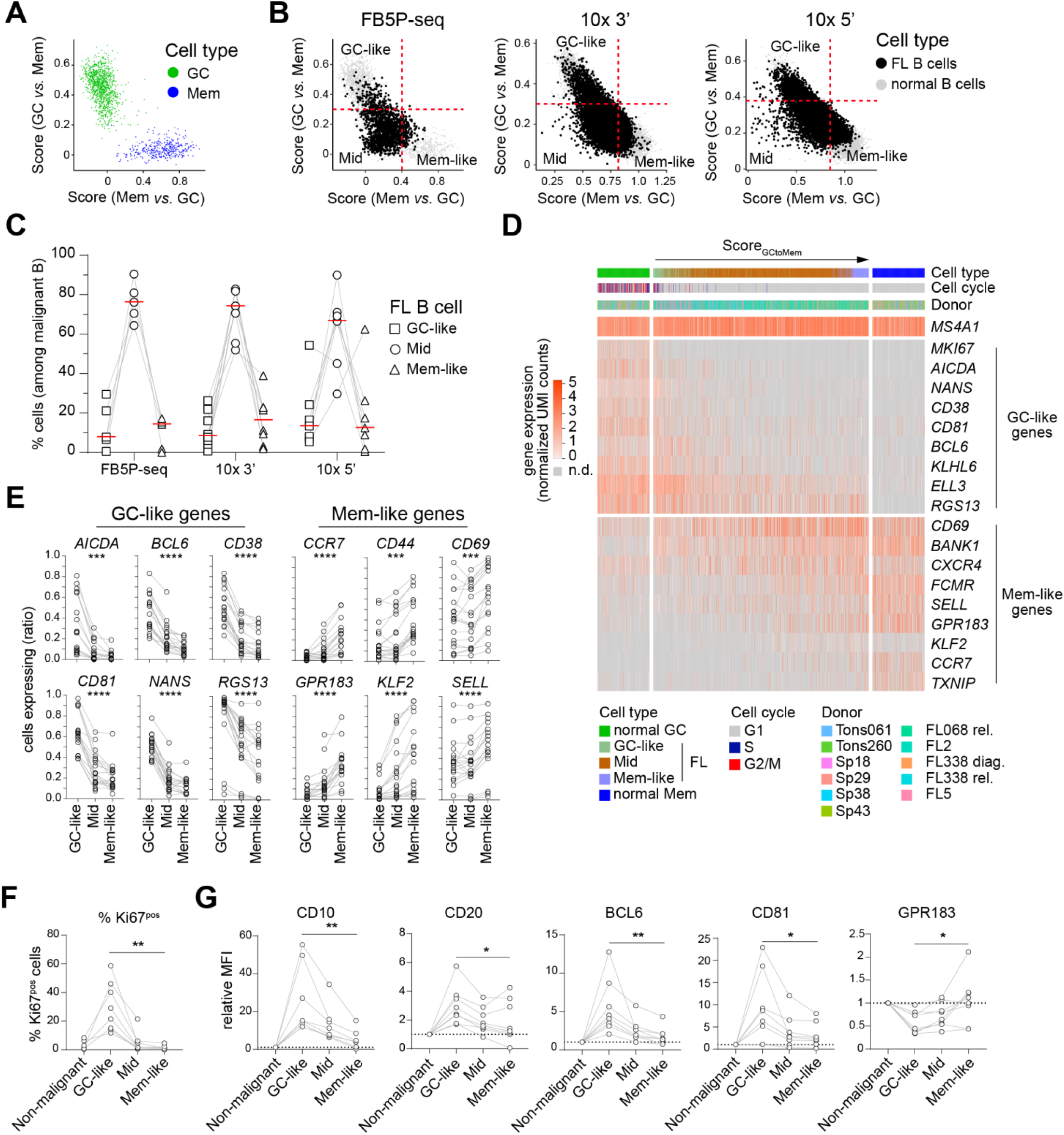
FL B cells adopt a continuum of transitional states between GC-like and Mem-like cell states. **(A)** GC *vs*. Mem signature score plotted against Mem *vs*. GC signature score for single normal spleen and tonsil GC and Mem B cells (FB5P-seq) colored by cell type, presented as a scatter plot. **(B)** GC *vs*. Mem signature score plotted against Mem *vs*. GC signature score for malignant FL B cells (black dots), presented as a scatter plot. Grey dots correspond to single normal spleen and tonsil B cells. Red dashed lines represent cut-offs used for discriminating GC-like (top left quadrant), Mid (bottom left quadrant) and Mem-like (bottom right quadrant) states. From left to right, data from FB5P-seq, 10x 3’ and 10x 5’. **(C)** Per-sample quantification of GC-like, Mid and Mem-like FL B cells for each scRNA-seq dataset. Quantification was calculated as the percentage of GC-like, Mid or Mem-like FL B cells among malignant FL B cells in each sample. Grey lines connect identical samples. *n*= 5 samples for FB5P-seq. *n*= 8 samples for 10x 3’. *n*= 7 samples for 10x 5’. **(D)** Gene expression heatmap for normal GC B cells (left), total FL B cells (middle) ordered by Score_GCtoMem_ values, and normal Mem B cells (right) analyzed by FB5P-seq. Cell type, cell cycle phase, and donor origin informations for each cell (columns) are color-coded in the bars above plot. Genes were selected on the basis of their expression in total B cells (*MS4A1*), GC B cells (next top half), or Mem B cells (bottom half). **(E)** Expression of GC-like (*AICDA, BCL6, CD38, CD81, NANS, RGS13*) and Mem-like (*CCR7, CD44, CD69, GPR183, KLF2, SELL*) genes in GC-like, Mid and Mem-like malignant FL B cells, computed as the ratio of cells expressing the indicated gene (expression >0) over the total cell number, for each cell subset of each sample. Black lines connect identical samples. *n*= 20 samples. ***p<0.001, ****p<0.0001, ns: not significant (Wilcoxon matched-pairs signed rank test comparing GC-like to Mem-like). **(F)** Percentage of Ki67 positive cells among non-malignant B cells and GC-like, Mid and Mem-like FL B cells. Grey lines connect identical samples. *n*= 8 samples. **(G)** Per-sample relative mean fluorescence intensities (MFI) for CD10, CD20, BCL6, CD81 and GPR183 in GC-like, Mid and Mem-like FL B cells normalized to non-malignant B cells. Grey lines connect identical samples. Black dotted lines represent baseline expression in non-malignant B cells. *n*= 8 samples. **(H-J)** *p<0.05, **p<0.01 (Mann-Whitney test, GC-like *vs*. Mem-like).

Unlike Mem-like cells, GC-like cells expressed high levels of *CD38* and *CD81* genes. We thus tested whether CD38 and CD81 surface markers could be used for enrichment of the two subsets by FACS. We sorted putative GC-like (CD38^hi^CD81^hi^) and Mem-like (CD38^neg^CD81^neg^) FL B cells from three human FL samples (**Fig.S4A-B**) and analyzed the cells with FB5P-seq. CD38^hi^CD81^hi^ cells expressed the GC-like transcriptional program, while CD38^neg^CD81^neg^ cells expressed Mem-like features (**Fig.S4C-D**). We then designed a flow cytometry analysis to analyze surface and intracellular protein markers in FL B cell subsets (**Fig.S1C**). GC-like cells displayed higher Ki67, CD10, CD20, BCL6, and CD81 protein expression levels (**Fig.3F-G**) and were larger in size (not shown), consistent with cell cycle activity. By contrast, Mem-like FL B cells were characterized by significantly increased expression of GPR183, an extra-follicular homing receptor (**Fig.3G**).

Thus, the vast majority of malignant B cells from the FL samples we studied spanned a continuum of cell states, ranging from proliferating GC-like to quiescent Mem-like cells, with identifiable phenotypic and functional features.

### Main source of intra-tumoral transcriptional heterogeneity

We then investigated intra-sample heterogeneity in each sample individually. We identified the GC-like to Mem-like gene expression continuum in every FL patient. In UMAP embeddings of sample-specific malignant B cells gene expression datasets, Score_GCtoMem_ values varied continuously from one end of the cloud of cells to another (**Fig.4A**). That observation suggested that the GC-like to Mem-like continuum was the main axis of transcriptional heterogeneity within FL B cells. In principal component analysis (PCA), the first principal component (PC1) cumulates the largest proportion of gene expression variance and represents the main axis of gene expression heterogeneity in a scRNA-seq dataset. In 13/15 samples, PC1 scores were tightly correlated with Score_GCtoMem_ values (**Fig.4B-C**). Unsupervised pseudo-time (PT) analysis, such as the one performed in the Monocle2 algorithm^43^, may identify cellular trajectories representing the most likely axes of variation (interpreted as differentiation axes) in a scRNA-seq dataset. When analyzing malignant B cells of each sample individually, PT ordering was also tightly correlated to the Score_GCtoMem_ continuum for 12/15 FL samples (**Fig.4D-E**). Together, PCA and unsupervised PT analyses clearly demonstrated that the GC-like to Mem-like continuum was the main source of intra-tumoral transcriptional heterogeneity in malignant B cells.

**Figure 4.**
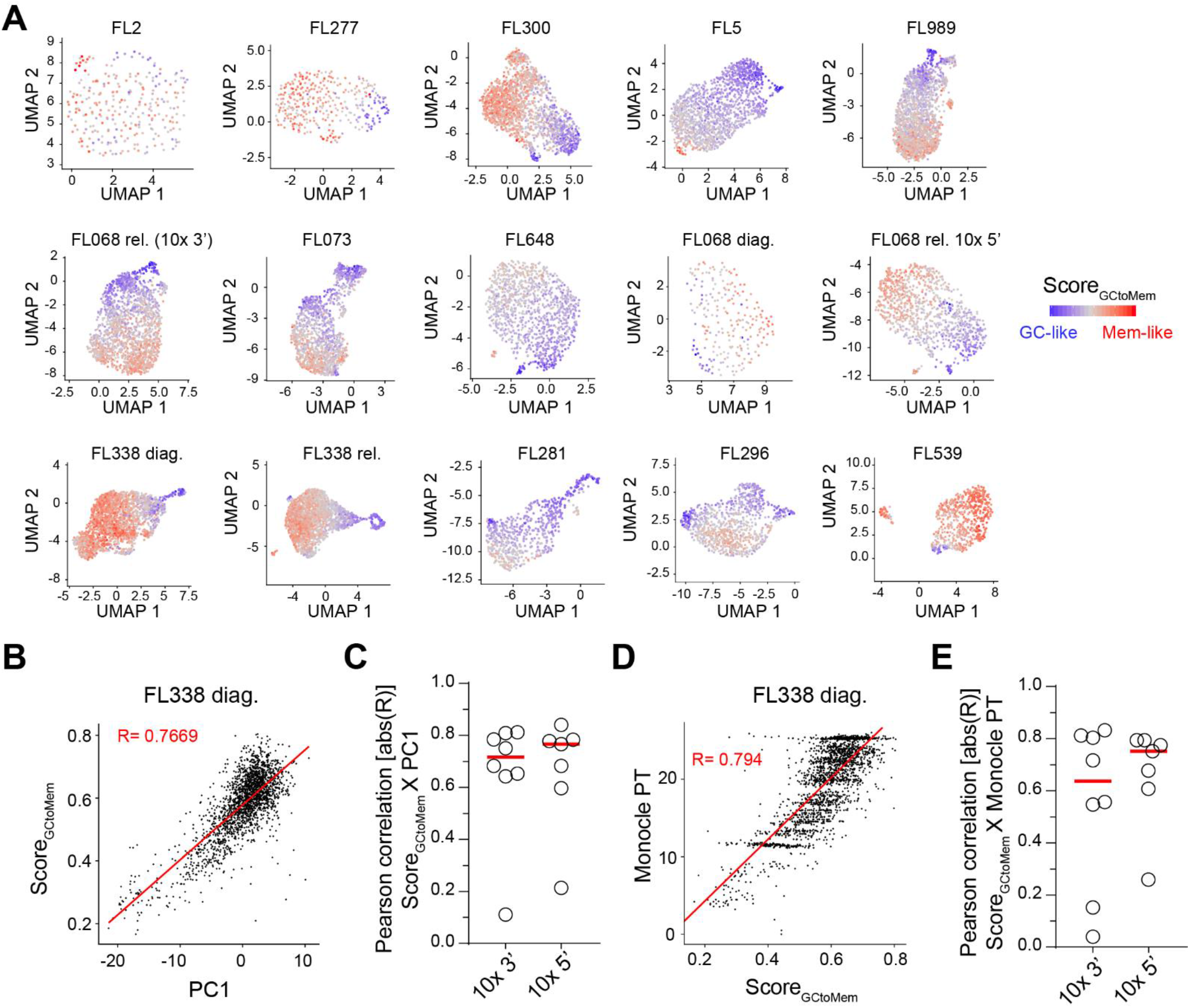
The GC-like to Mem-like continuum is the main source of intra-tumoral transcriptional heterogeneity in FL B cells. **(A)** Per-sample UMAP embeddings of total FL B cells from 10x 3’ (FL2, n=269; FL277, n=352; FL300, n=1748; FL5, n=1421; FL989, n=1798; FL068 rel., n=1740; FL073, n=1354; FL648, n=1136), or 10x 5’ (FL068 diag., n=225; FL068 rel., n=968; FL338 diag., n=2133; FL338 rel., n=2127; FL281, n=638; FL296, n=897; FL539, n=621) scRNA-seq datasets. Each symbol represents an individual cell. Cells are colored according to their Score_GCtoMem_ value, from the most GC-like (blue) to the most Mem-like (red). **(B)** Scatter plot of Score_GCtoMem_ *vs*. PC1 score values for FL B cells from the FL338 diag. sample analyzed by 10x 5’. Red line represents linear regression. R: Pearson correlation coefficient. Each symbol represents an individual cell. **(C)** Per-sample Pearson correlation values between Score_GCtoMem_ and PC1 score for total FL B cells from samples analyzed by 10x 3’ (n=8) or 10x 5’ (n=7) scRNA-seq as indicated. **(D)** Scatter plot of Monocle-defined pseudo-time score (Monole PT) *vs*. Score_GCtoMem_ values for FL B cells from the FL338 diag. sample analyzed by 10x 5’. Red line represents linear regression. R: Pearson correlation coefficient. Each symbol represents an individual cell. **(E)** Per-sample Pearson correlation values between Score_GCtoMem_ and Monocle PT for total FL B cells from samples analyzed by 10x 3’ (n=8) or 10x 5’ (n=7) scRNA-seq as indicated.

### Gene expression states are independent of subclonal BCR evolution

Malignant B cell gene expression heterogeneity may stem from divergent subclonal genetic evolution or cell plasticity. Through the activation or the deletion of transcriptional regulators, genetic divergence may lead to transcriptional changes in subclones which will then appear as transcriptionally distinct cell states. Alternatively, cell plasticity, *i*.*e*. the ability of cells to dynamically shift from a transcriptional state to another, could produce a continuum of different cell states. We took advantage of subclonal BCR evolution as a proxy to track genetic evolution. In FL, ongoing somatic hypermutation (SHM) of the *IGH* and *IGK/L* genes^13^ facilitates the identification of malignant subclones to track long term subclonal evolution^44^. Through single-cell analysis of paired *IGH* and *IGK/L* sequences from 3912 FL B cells, we tracked FL B cell subclonal evolution for 7 FL tumors (**Fig.5, S5, Supplementary Data Table 4**).

**Figure 5.**
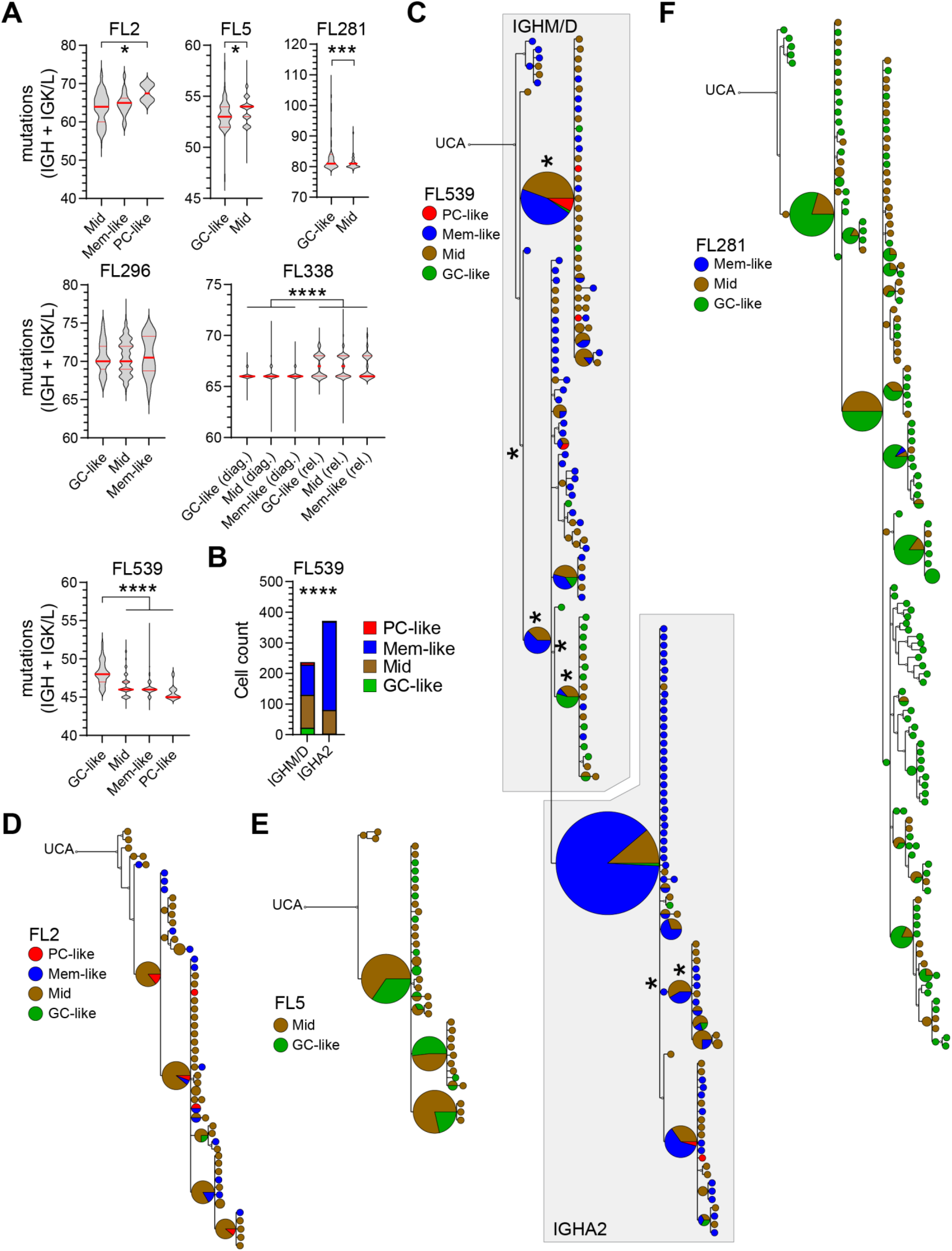
FL B cell gene expression states are independent of subclonal BCR divergence. **(A)** Violin plots representing the distributions of total number of somatic mutations in *IGH* and *IGK/L* genes for FL B cells of the indicated subsets (GC-like, Mid, Mem-like, PC-like) and samples (FL2, FL5, FL281, FL296, FL338, FL539) as compared to the sample-specific umutated common ancestor sequence. Bold and thin red lines indicate median and interquartile range, respectively. *p<0.05, ***p<0.001, ****p<0.0001 (Mann-Whitney or Kruskal-Wallis test). **(B)** Bar plot representing the number of FL B cells from the indicated subset (color scheme, GC-like, Mid, Mem-like, PC-like) of FL539 expressing either an *IGHM/D* or an *IGHA2* isotype. ****p<0.0001 (Chi-square test). **(C-F)** Phylogenetic trees built from concatenated *IGH*-*IGK/L* nucleotide sequences of clonally related single malignant FL B cells for samples FL539 **(C)**, FL2 **(D)**, FL5 **(E)** and FL281 **(F)**. The non-filled circle symbol (top left) indicates the inferred UCA sequence (unmutated common ancestor) to which the tree is rooted. Each symbol in the tree corresponds to a unique *IGH-IGK/L* sequence; symbol size is relative to the number of cells carrying that sequence. Symbols are colored according to cell subset (PC-like: red, GC-like: green, Mem-like: blue, Mid: brown). In the case of multiple cells carrying the same sequence, proportions of cells in each state are indicated as a pie chart within the symbol. For FL539, the subclones expressing either an *IGHM/D* or an *IGHA2* isotype are circled by grey boxes. * besides a node indicates a statistically significant difference in cell states (colors) proportions among cells below that node compared to the whole sample.

In FL281 and FL539, GC-like cells were significantly more mutated than other subsets (**Fig.5A**), consistent with GC-like cells actively cycling and expressing *AICDA* required for ongoing SHM (**Fig.3D-F**). Analysis of FL338 paired diagnosis/relapse samples showed the emergence of a relapse-specific subclone with more mutations (**Fig.5A, S5A-B**). In FL539, we detected two subclones expressing either IGHM/D or IGHA2 isotype that differed in state composition (**Fig.5B-C**). Phylogenetic trees built from the single-cell BCR sequences (**Fig.5C-F, S5**) showed that cells in PC-like, GC-like, Mid and Mem-like states were randomly distributed across BCR subclonal branches in 6/7 samples. Only sample FL539 showed a statistically significant imbalance, with some subclonal branches dominated by Mid or GC-like states (**Fig.5C**), although other cell states were also present in those BCR subclones. Those data argue against subclonal genetics directly imprinting main FL B cell transcriptional states, and support the hypothesis that PC-like, GC-like, Mid and Mem-like states are the result of functional plasticity.

### Impact of T_FH_-derived activation signals

Interactions with TME may drive FL B cell plasticity. Non-supervised clustering of TME cells from the integrated 10x 3’ and 10x 5’ datasets defined 15 distinct clusters (**Fig.6A, S6A**). Parallel analysis of TME cell types by flow cytometry (**Fig.S1D**) demonstrated tight correlations between cell subset frequencies in scRNA-seq and FACS (**Fig.S6B**). The TME composition was variable and characterized by the predominance of CD4^+^ T cells, with high frequencies of T central memory (T_CM_), T_FH_ and T regulatory (T_REG_) subsets (**Fig.6B, S6C**). T_FH_ cells were the main producers of *CD40LG, IL4* and *IL21* mRNA (**Fig.6C, S6D**), encoding critical positive regulators of B cell activation, proliferation and differentiation.

**Figure 6.**
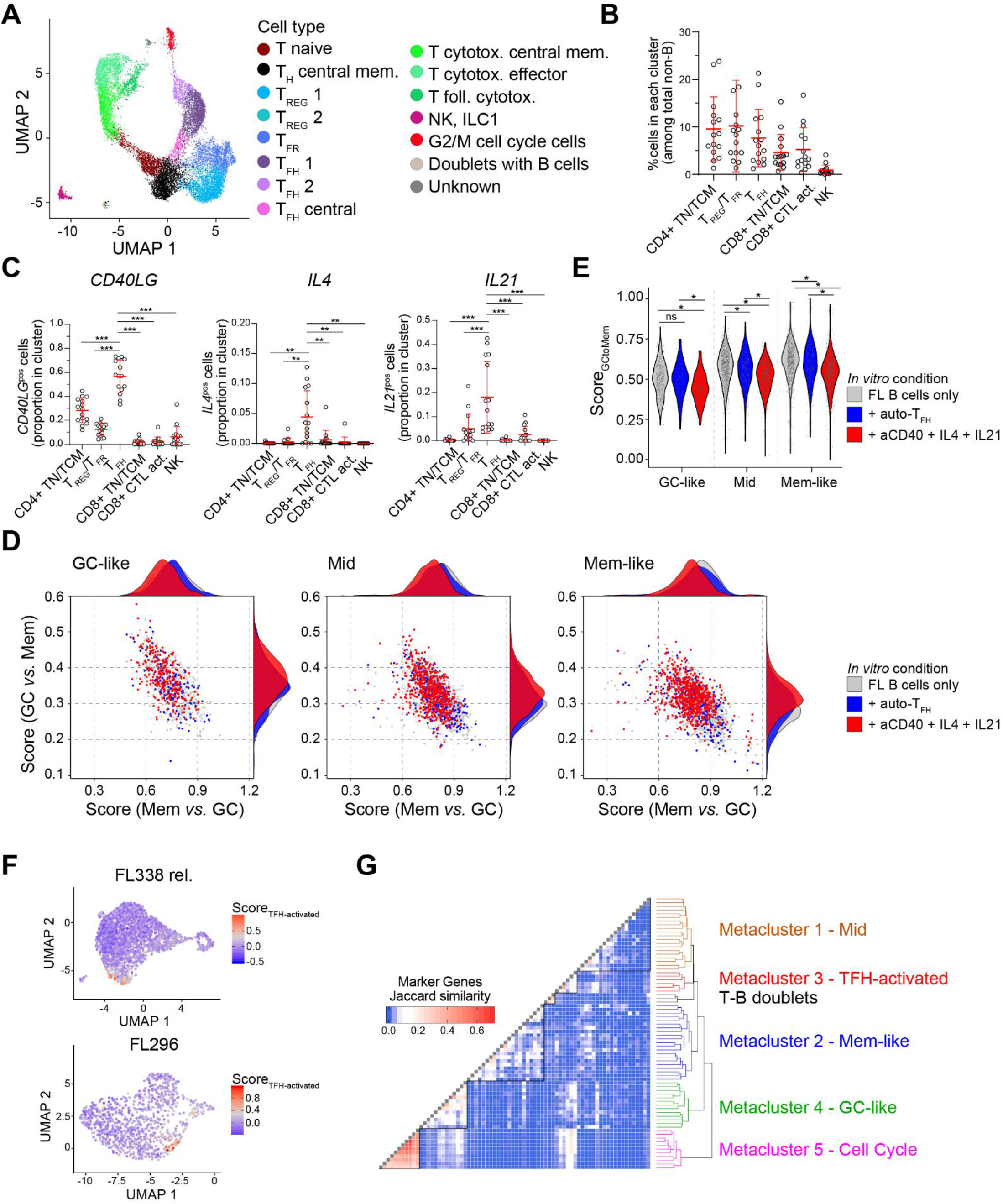
T_FH_-derived activation signals impact FL B cell states *in vitro* and *in vivo*. **(A)** UMAP embedding of single TME non-B cells from FL cell suspensions analyzed by 10x 3’ and 10x 5’ scRNA-seq as described in **Fig.1**, after integration. Cells are colored by cell types defined by unsupervised clustering and manual annotation. **(B)** Per-sample quantification of CD4+ naive/central memory T cells (CD4+ TN/TCM), T regulatory/T follicular regulatory cells (T_REG_/T_FR_), T follicular helper cells (T_FH_), CD8+ naive/central memory T cells (CD8+ TN/TCM), CD8+ activated cytotoxic T cells (CD8+ CTL act.) and Natural killer cells (NK). Quantification was calculated as the percentage of cells found in each cell subset (identified in **(A)**, some clusters grouped as described in Methods) among total TME non-B cells for each sample. *n*= 15 samples. Red bars indicate mean and standard deviation. **(C)** Per-sample quantification of cells expressing *CD40LG* (left), *IL4* (middle), or *IL21* (right), among cell types defined in **(B)**. Quantification was calculated for each sample as the ratio between the number of cells with non-zero expression of the indicated gene in each subset and the total cell number in that subset. *n*= 15 samples. **p<0,01 ***p<0.001 (Wilcoxon matched-pairs signed rank test with Bonferroni correction). Red bars indicate mean and standard deviation. **(D)** Scatter plots and marginal densities of Mem *vs*. GC (x-axis) and GC *vs*. Mem (y-axis) gene expression scores for FL B cells from the indicated subsets (GC-like: left, Mid: middle, Mem-like: right) after 48-hour *in vitro* culture in the indicated conditions: FL B cells only (grey), in presence of 1:1 ratio of autologous T_FH_ cells (+ auto-T_FH_, blue), or in presence of T_FH_-derived activation signals (+ aCD40 + IL4 + IL21, red). Cells from FL068 diagnosis/relapse and FL338 diagnosis/relapse samples were analyzed separately but pooled in the analysis. Each dot represents a single cell. **(E)** Violin plot and dot plot of Score_GCtoMem_ values for sorted GC-like (left), Mid (middle) and Mem-like (right) FL B cells after 48-hour *in vitro* culture in the indicated conditions (same color code as **(D)**). Cells from FL068 diagnosis/relapse and FL338 diagnosis/relapse samples were analyzed separately but pooled in the analysis. Each dot represents a single cell. *p<0.05, ns: not significant (Wilcoxon rank sum test with Benjamini & Hochberg adjustment). **(F)** Per-sample UMAP embeddings of total FL B cells from FL338 rel. (top) and FL296 (bottom) samples. Each symbol represents an individual cell. Cells are colored according to their Score_TFH-activated_ value. **(G)** Hierarchical clustering of pairwise marker genes Jaccard similarity matrix of 74 sample-specific graph-based malignant FL B cell clusters from 15 scRNA-seq datasets, highlighting metacluster 1 (Mid, brown), metacluster 3 (T_FH_-activated, red), T-B doublets metacluster (black), metacluster 2 (Mem-like, blue), metacluster 4 (GC-like, green) and metacluster 5 (Cell Cycle, magenta).

T_FH_ cells control normal B cell differentiation at multiple stages of GC responses^45^. We hypothesized that FL T_FH_ cells could trigger malignant B cell plasticity. We sorted GC-like, Mid and Mem-like malignant B cells from 4 FL samples (**Fig.S7A**) and cultured them *in vitro* for 48h, either alone (FL B cells only), with autologous T_FH_ cells (**Fig.S7B**), or with T_FH_-mimicking signals (anti-CD40 activating antibody, IL-4 and IL-21). After barcoding, we analyzed live cells in all culture conditions by 10x 5’ scRNA-seq. In the absence of external stimuli, FL B cell subsets conserved Score_GCtoMem_ values consistent with their sorting phenotype (**Fig.6D-E, S8A**). In the presence of T_FH_ or T_FH_-mimicking signals, both Mem-like and Mid cells showed a decrease of Score_GCtoMem_ values indicating transcriptional change towards a GC-like state (**Fig.6D-E, S8A**). The decrease in Score_GCtoMem_ was stronger with T_FH_-mimicking signals than with autologous T_FH_ cells. GC-like cells cultured with T_FH_-mimicking signals, but not autologous T_FH_ cells, also showed a lower Score_GCtoMem_. Thus, T_FH_-derived signals impacted B cell transcriptional states *in vitro*, “pushing” malignant B cells towards a GC-like state.

To track whether such dynamics also occurred *in vivo*, we defined a set of 25 genes whose expression was robustly increased in malignant B cells upon co-culture with autologous T_FH_ *in vitro* (**Fig.S8B-C**). We computed the corresponding signature score (Score_TFH-activated_) in malignant B cells from 10x 3’ and 10x 5’ datasets. In 10/15 samples, a minor subset of malignant B cells with high Score_TFH-activated_ values clustered together in sample-specific UMAP (**Fig.6F**), suggesting those cells had been recently activated by T_FH_ cells *in vivo*. Unsupervised marker genes-based metaclustering (detailed in **Computational Methods**) of the 74 sample-specific malignant B cell clusters from the 15 scRNA-seq datasets identified 6 distinct metaclusters (**Fig.6G**). Analysis of shared marker genes (**Supplementary Data Table 5**) and single-cell signature scores (**Fig.S8D-F**) allowed to annotate those unsupervised metaclusters as GC-like, Mid, Mem-like, CellCycle, and T_FH_-activated. Cells from the T_FH_-activated metacluster expressed intermediate levels of Score_GCtoMem_ (**Fig.S8D**) but high levels of Score_TFH-activated_ (**Fig.S8F**), suggesting they corresponded to Mid cells with a specific activated profile.

Thus T_FH_-derived signals trigger a conserved T_FH_-activated state in a minor subset of malignant B cells *in vivo*. T_FH_-activated cells are distinct from Mem-like or GC-like cells.

### Localization of malignant B cell subsets

In lymph nodes, GC and Mem B cells home to central and more peripheral areas of B cell follicles, respectively^46^. FL tumors are often structured as adjacent or coalescent B cell follicles which contain malignant B cells^47^, but a variable proportion of malignant B cells are present also in interfollicular zones^48^. As several of the marker genes of Mem-like B cells encoded receptors involved in perifollicular or extrafollicular homing (*CCR7, CXCR4, GPR183*), we hypothesized that distinct malignant B cell states may localize to specific areas of FL lymph nodes.

We performed spatial transcriptomics analysis (ST) of a lymph node section from a grade 1-2 FL tumor. We focused our analysis on the region that contained tumor follicles (**Fig.7A**), and defined distinct tissue areas relative to follicles (**Fig.7B, S9A**): centrofollicular (CF), perifollicular (PF), interfollicular (IF) and extrafollicular (EF). ST slides capture mRNA on 55-µm diameter spots, thereby averaging the transcriptome of several cells (5 to 20 cells in lymphoid tissues). We constructed a reference scRNA-seq dataset of FL cells by integrating our full 10x 3’ and 10x 5’ datasets (**Fig.7C**), and applied reference-based deconvolution to infer the presence of malignant and non-malignant cell subsets in each spot^49^. The deconvolution algorithm *cell2location* outputs q05_spot_factors, probabilistic values reflecting the number of cells of a given cell type in ST spots^49^. For most of the cell types/states assessed, there was a moderate to good correlation between q05_spot_factors and signature scores based on marker genes (**Fig.S9B**, R>0.4 for 10/15 cell types), confirming that the deconvolution effectively localized cell types/states *in situ*. The sum of q05_spot_factors, reflecting the total number of cells per spot, was highest in CF spots and gradually decreased from CF/PF to IF/EF areas (**Fig.S9C**), consistent with higher cell densities in CF areas.

**Figure 7.**
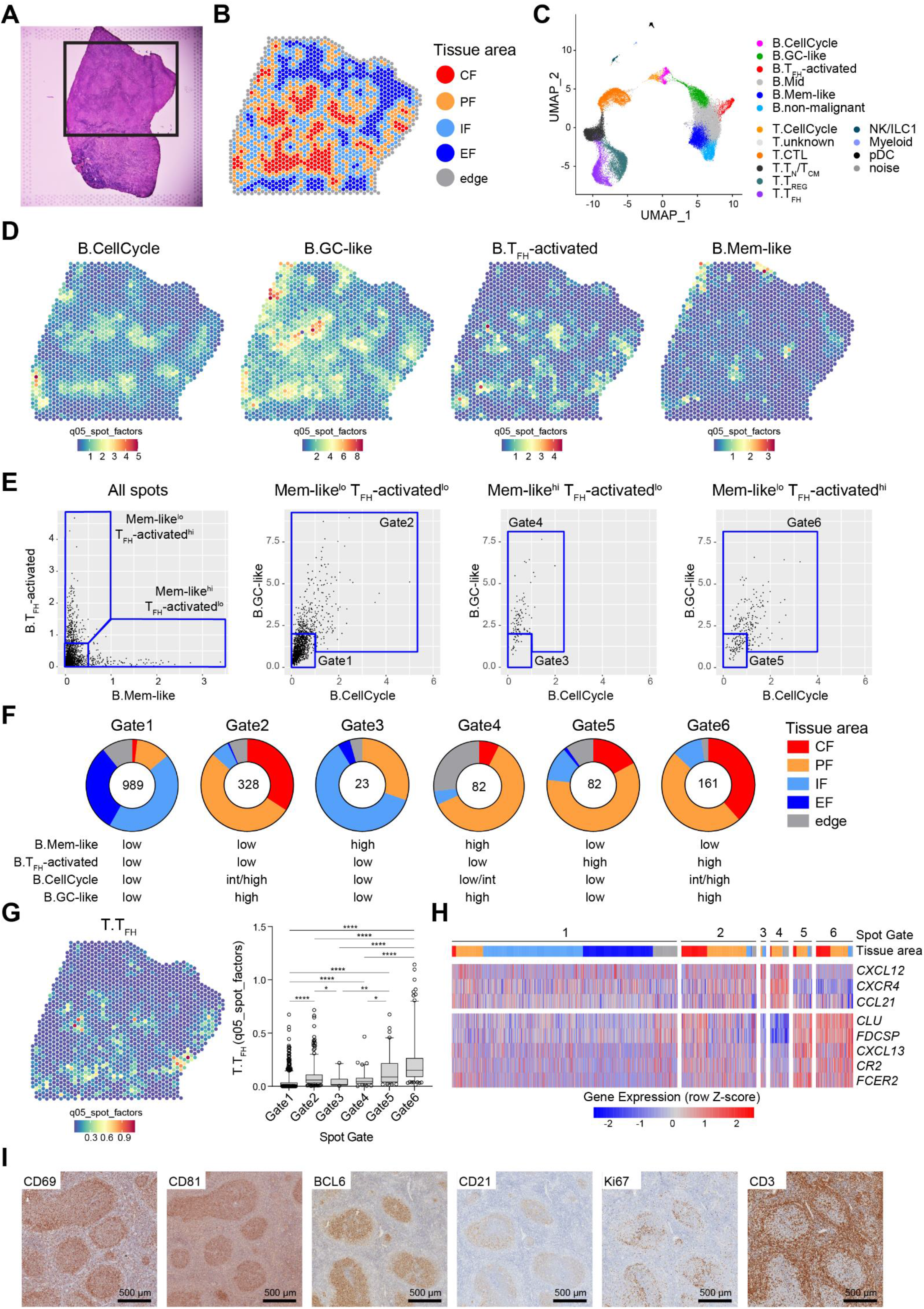
Spatial distribution of malignant B cell subsets in FL lymph node. **(A)** Hematoxylin and Eosin stained image of FL lymph node analyzed by spatial transcriptomics. Black square: region of interest containing tumor follicles. **(B)** Spatial transcriptomics spot map colored by tissue area: CF centrofollicular, PF perifollicular, IF interfollicular, EF extrafollicular. **(C)** UMAP embedding of all single live cells from FL cell suspensions analyzed by 10x 3’ and 10x 5’ scRNA-seq after integration. Cells are colored by cell types defined by unsupervised clustering and manual annotation. **(D)** Spatial transcriptomics spot maps, colored by q05_spot_factors values for the indicated cell subsets after deconvolution with *cell2location*. **(E)** Gating strategy of spatial transcriptomics spots based on malignant B cell content after deconvolution. Scatter plots show q05_spot_factors of the indicated cell types. Each dot is a spatial transcriptomics spot, gated as indicated by the blue quadrants. Leftmost plot shows the first gating layer on all spots, then from left to right, the subgating of Mem-like_lo_ T_FH_-activated^lo^, Mem-like^hi^ T_FH_-activated^lo^, and Mem-like^lo^ T_FH_-activated^hi^ spots, respectively. **(F)** Pie charts of tissue area types, as defined in (B) for spatial transcriptomics spots gated in (E). Numbers within each pie chart indicate the total number of spots in the gate. **(G)** Spatial transcriptomics spot map, colored by q05_spot_factors values for the T.TFH cell subset (left), and box plot of T.TFH q05_spot_factors values for spots gated in (E).*p<0.05, **p<0.01, **** p<0.0001, Kruskal-Wallis test with multiple comparisons and Dunn’s correction. **(H)** Row-normalized (z-score) gene expression heatmap for spatial transcriptomics spots grouped by gates shown in (E), for the indicated genes. Tissue area type (B) of each spot is color-coded on the panel above the heatmap. **(I)** Immunohistochemistry detection of CD69, CD81, BCL6, CD21, Ki67 and CD3 proteins in sections from FL073 lymph node biopsy sample (Supplementary Tables 2 and 3). Scale bars = 500μm.

Malignant B cell states mapped to different areas of the tumor (**Fig.7D**). We classified spots based on the inferred densities of distinct malignant B cell states (gates in **Fig.7E**), and quantified the area type distribution for each spot gate (**Fig.7F**). GC-like and Mem-like states tended to segregate in CF and IF areas, respectively. Multiple states were mixed in PF areas, which were also where T_FH_-activated states seemed most abundant. T_FH_ q05_spot_factors were significantly higher in spots containing T_FH_-activated B cells (**Fig.7G**), suggesting activation of malignant B cells with T_FH_ occurred mostly in PF areas. Stromal cells play important roles in lymphoid cell localization and function in normal and malignant lymph nodes^30^, but those cells were absent from our scRNA-seq reference for deconvolution. Nevertheless, Mem-like^hi^ spots expressed higher levels of *CXCL12* and *CCL21*, encoding T-zone reticular cell-derived chemo-attracting ligands of Mem-like receptors *CXCR4* and *CCR7*, respectively (**Fig.7H**). By contrast, markers of B-zone follicular dendritic cells (FDC) were detected at higher levels in T_FH_-activated^hi^ spots (**Fig.7H**), suggesting that distinct stromal cell types provide survival and/or activation niches for distinct malignant B cell states. Although our ST analysis was performed on only one FL sample, single-color immunohistochemistry staining of markers of Mem-like cells (CD69, staining mostly PF and IF areas), GC-like cells (CD81, BCL6, NANS, CD38, staining CF areas), FDC (CD21, staining outer CF and PF areas), proliferating cells (Ki67, staining CF and PF areas) and T cells (CD3, CD5, staining mostly PF and IF areas) in tissue sections from six FL samples (**Fig.7I, S10**) were consistent with the localizations inferred from ST.

## DISCUSSION

Tumors are dynamic ecosystems where malignant and non-malignant cell types co-exist. Here we have combined supervised and non-supervised analyses of integrative single-cell transcriptomics datasets to uncover the conserved states of malignant B cells in FL. Beyond inter-patient divergence^37,38^, our work reveals five recurrent malignant B cell states: GC-like, Mid, Mem-like, T_FH_-activated and PC-like states. Our analyses support a model where FL B cell states span a continuum from quiescent inter- and peri-follicular Mem-like to proliferative centro-follicular GC-like state, with most malignant B cells being in a quiescent non-activated Mid state within follicles. In that model, activation of Mem-like and Mid cells through CD40-dependent interactions with T_FH_ cells, which may occur in peri-follicular FDC-rich areas, induces a specific T_FH_-activated state which likely serves as an intermediate towards the GC-like state. That model fits with previous observations, provides a framework for understanding FL B cell biology, and may be useful for designing novel therapies.

So far, only two subsets of FL B cells had been identified^50,51^: FL centrocytes and centroblasts, named based on morphological similarities with GC B cell types. Those cell subsets have been the basis of the histologic grading system of FL for decades^52^. Our work refines and updates FL centroblast / centrocyte discrimination by characterizing previously unrecognized cell states. The centrocyte and centroblast names are misleading, because only GC-like cells, which most likely correspond to centroblasts, are transcriptionally and functionally close to normal GC B cells. By contrast, non-cycling Mem-like, Mid and T_FH_-activated cells, which encompass cells that would be considered as FL centrocytes, express non-GC transcriptional and functional programs. In fact, the CXCR4 and CD83 markers which segregate normal GC centroblasts (CD83^lo^CXCR4^hi^) from centrocytes (CD83^hi^CXCR4^lo^)^35^, are markers of Mem-like (*CXCR4*) and T_FH_-activated (*CD83*) FL B cell states, and should not be used to discriminate FL centroblasts and centrocytes.

We define Mem-like B cells as small quiescent lymphocytes expressing genes and proteins associated with peri- or inter-follicular homing (*GPR183, CCR7, CXCR4*). Our *in situ* analyses showed that Mem-like cells are located in peri-follicular and inter-follicular areas and are not found within tumor follicles. The Mem-like state thus likely corresponds to the interfollicular FL B cells which have been reported and studied by *in situ* analysis, laser microdissection and BCR sequencing^48,50,53^. Importantly, we detected those cells in all FL samples we studied, regardless of disease grade (grades 1-2 or 3a) and stage (diagnosis or relapse), suggesting that interfollicular circulation of malignant B cells is a universal feature that may contribute to the dissemination of disease at all stages of progression. In normal immune responses, Mem B cell reactivation triggers mainly differentiation into PC, but some cells may re-enter GC reactions in T_FH_-dependent manner^5,7,54^. Mouse models have shown that BCL2-overexpressing Mem B cells favor the GC fate when reactivated^26^. Our *in vitro* experiments showed that Mem-like cell activation with autologous T_FH_ or T_FH_-mimicking conditions initiates GC-like differentiation, but very limited and non-significant PC-like differentiation (not shown). Those data suggest that Mem-like cells are poised to differentiate to GC-like cells when activated by T_FH_ *in vivo*, and only very rarely initiate plasmacytic differentiation.

Nonetheless, we detected PC-like cells by supervised analysis in 7/15 samples. PC-like cells were detected in more samples when using droplet-based 10x 3’ and 10x 5’ methods, which is likely due to the higher throughput of those methods compared to FB5P-seq, enabling to more frequently capture those rare cell types. Those cells were in sufficient numbers to be properly characterized only in the samples from two patients (FL2 and FL539, 2-3% of malignant cells). Those patients may correspond to cases of FL with plasmacytic differentiation^41^. PC-like cells express all features associated with high levels of active antibody production, but do not express the mature plasma cell marker *SDC1* (encoding CD138), suggesting that plasmacytic differentiation of malignant B cells is incomplete. In the future, higher throughput single-cell analyses of larger cohorts of FL patients will help to decipher the signals involved in PC-like differentiation and whether some mutational landscapes might be more permissive to plasmacytic differentiation.

The T_FH_-activated state is characterized by expression of adhesion (*ICAM1*), activation (*CD83, CD40*) and CD40 response genes (*EBI3, CCL17, MIR155-HG, BATF*), some of which were already shown to be induced by FL B:T_FH_ co-cultures^32^ or in activated light zone GC B cells^55^. The supportive pro-survival role of T_FH_ cells in FL tumors, notably through CD40L and IL4 signals, has been described previously^31,32,56,57^, but whether all malignant B cells were equally supported was not known. We show that T_FH_-activated cells are only a minor fraction of malignant FL B cells at any time, and may be located mostly in perifollicular areas where strong FDC-derived mRNA expression is also detected. This suggests that most productive interactions between malignant B cells and supportive T_FH_ cells may occur in specific niches at the periphery of tumor follicles.

We provide evidence that malignant B cells are plastic and dynamically alternate between GC-like and Mem-like states *in vivo*. The SHM process of BCR mutation requires AID and cell cycling^16,58,59^, two exclusive features of the GC-like state. If transitions from GC-like to Mid and Mem-like were rare, novel BCR mutations would ‘preferentially’ accumulate in GC-like cells, leading to single-cell BCR phylogenetic trees where GC-like cells would accumulate at the most terminal leaves. While that may be the case in some samples (*e*.*g*. FL281), in most samples, cells from any given BCR subclone were found in different states, even in tumors with extensive ongoing SHM. Thus, we gather that in the majority of FL tumors, plasticity occurs more frequently than BCR SHM. By analogy with normal B cell dynamics^46^, cell state transitions along the neoplastic GC-Mem continuum may occur on a timescale of one to a few days.

In most samples, the majority of malignant B cells are in Mid state, where cells express rather low levels of either GC-like or Mem-like genes. Remarkably, Mid cells were the least conserved state, with almost no markers of those cells being shared among multiple patients. This suggests that Mid cells represent the default state of equilibrium of malignant B cells, which is enforced by patient-specific genetic abnormalities. By contrast, the GC-like, Mem-like, T_FH_-activated and PC-like states reflect interactions between malignant B cells and their microenvironment, which represent conserved signatures of FL biology. Those conserved states likely underlie various biological processes such as progression (GC-like) and dissemination (Mem-like). The frequency of distinct malignant states among FL B cells is variable from sample to sample. Those different compositions may be due to different genetics, and/or different TME composition. B cell lymphomas often present as disseminated diseases with multiple lymph nodes involved^11^. Our analyses clearly show that the most conserved FL states (GC-like, Mid, Mem-like, T_FH_-activated) can be observed in all FL samples and reflect transcriptional dynamics occurring in parallel in multiple tumor follicles. Yet it is unclear to what extent tumor composition reflects the general state of a patient’s tumor, or may be subject to variations from site to site. Distinct sites may harbor genetically distinct malignant B cell subclones^60^. In a study exploring site-to-site heterogeneity in FL, integrative scRNA-seq revealed that malignant B cells in separate lymph nodes, sampled from the same patient at a given time, expressed divergent transcriptional programs in proportions commensurate to genetic divergence assessed by subclonal BCR diversity^61^. TME composition was more consistent across sites, but the extent of T_FH_ infiltration varied and those variations were correlated to site-to-site malignant B cell dissimilarity. Altogether, those data suggest that the frequencies of conserved malignant B cell states among a tumor lymph node biopsy will be subject to site-to-site variations. Future scRNA-seq studies on larger cohorts will inform whether tumor composition in conserved malignant states can be predictive of clinical course, beyond statistical noise due to site-to-site variations.

Distinct biological states within a tumor may exhibit distinct sensitivities to therapy, and/or distinct abilities to evade therapy and seed relapse. A study using scRNA-seq on B cell lymphomas demonstrated divergent drug sensitivities of subclones within a single tumor^38^. Standard first-line treatment for FL combines anti-CD20 antibodies and chemotherapy^11^. GC-like cells are actively proliferating and express high levels of CD20, and therefore may be efficiently targeted by current treatment options. Mem-like B cells are quiescent and express lower levels of CD20, therefore those cells may be more prone to resist or evade current treatments. Moreover, the expression of extra-follicular homing markers in Mem-like cells likely confers a strong potential for dissemination, as previously shown for CXCR4, CD44 or CCR7 in other B cell neoplasia^62–65^. We speculate that Mem-like cells could reinitiate lymphoma if assisted by the appropriate T cell help, thereby acting as relapse initiating cells, a cell entity that has been postulated from genetic studies but whose actual identity has been elusive^66–68^. PC-like cells are also quiescent and express no CD20, which would make them resistant to first-line therapy, but reinitiating lymphomas from PC-like cells would imply to de-differentiate from the antibody-producing cell phenotype, which has never been observed in non-malignant PC. Given the functional plasticity we describe here, we anticipate that therapeutic strategies that do not target all malignant cell types in follicular lymphoma, including minor quiescent subsets, will fail to induce long remissions without relapses. Targeting the B-TME interactions that sustain malignant B cell plasticity is a promising avenue^69,70^.

## EXPERIMENTAL METHODS

### Human samples

Non-malignant spleen samples were obtained as previously described^26^ (**Table S1**). Non-malignant tonsil samples (**Table S1**) and most FL samples (**Table S2**) were obtained from the CeVi collection of the Carnot Calym Institute (ANR, France, *https://lymphoma-research-experts.org/calym/explore-the-resources-of-the-consortium/cevi-collection/)*. FL2 and FL5 sample biopsies were obtained from the cell bank of Hospices Civils de Lyon hospital (Centre de Ressources Biologiques des Hospices Civils de Lyon, **Table S2**) and were previously described^36^. All samples were preserved as frozen live mononuclear cell suspensions in liquid nitrogen or at -150°C until use. FFPE and frozen samples for immunohistochemistry and spatial transcriptomics were obtained from Hôpital Henri Mondor (**Table S2**). Written informed consent was obtained from the patients, donors or relatives in accordance with the Declaration of Helsinki and with IRB approval of the French Biomedicine Agency.

### Flow cytometry

Frozen spleen, tonsil or FL cell suspensions were thawed at 37°C in RPMI + 10% FCS, then washed and resuspended in FACS buffer (PBS + 5% FCS + 2mM EDTA) at a 10^8^ cells/ml concentration for staining. Cells were first incubated with 2% normal mouse serum (Life technologies) and Fc-Block (BD Biosciences or Biolegend) for 10 min on ice. Then cells were stained with mixes of fluorophore-conjugated antibodies to surface antigens (**Table S3**) for 30 min on ice. Cells were washed in PBS, then incubated with the Live/Dead Fixable Aqua Dead Cell Stain (Thermofisher) for 10 min on ice. If necessary, intracellular staining was performed after LiveDead staining. Cells were fixed and permeabilized using the Foxp3/Transcription Factor Staining Buffer set (Life Technologies) according to the manufacturer’s instructions. Cells were then stained with intracellular marker fluorophore-conjugated antibodies (BCL6 and Ki67, **Table S3**) for 30 min on ice. After a final wash in FACS Buffer, cells were resuspended in FACS Buffer and samples were acquired on a BD LSR Fortessa or a BD FACS Influx. Gating strategies for analyses and cell sorting are described in **Fig.S1, S4, S7**. Flow cytometry data were analyzed with FlowJo software.

### FB5P-seq

For the experiment described in **Fig.1A**, Mem B cells were gated as CD3- CD14- IgD- CD20+ CD10- CD38^lo^ CD27+ SSC^lo^ single live cells. GC B cells were gated as CD3- CD14- IgD- CD20+ CD10+ CD38^int^ single live cells. Besides sorting total GC B cells, we also focused cell sorting on CXCR4+ CD83+ GC B cells in some 96-well plates. For the FB5P-seq bioinformatics analysis, GC B cells were defined as the sorted total GC as well as the CXCR4+ CD83+ GC B cells. PC cells were gated as CD3- CD14- IgD- CD20- CD38^hi^ CD27+ SSC^hi^ single live cells. Early PC cells were gated as CD3- CD14- IgD- CD20+ CD38^hi^ CD27+ CD10+ single live cells (**Fig.S1B**). For FB5P-seq data analysis, PC cells were defined as combining PC and early PC cells.

For the experiment described in **Fig.1B**, FL B cells were gated as CD3- CD14- CD20+ CD10+ and on the basis of λ or κ light chain isotype restriction (**Fig.S1A**).

For the experiment described in **Fig.S4**, putative malignant GC-like cells were sorted as CD3- CD14- IgD- CD38^hi^ CD81^hi^ single live B cells and based on light chain isotype restriction. Putative malignant Mem-like cells were sorted as CD3- CD14- IgD- CD38^lo^ CD81^lo^ single live B cells and based on light chain isotype restriction. Total malignant CD3- CD14- IgD- single live B cells were also sorted based on light chain isotype restriction.

FB5P-seq protocol was performed as previously described^39^. Briefly, single normal or FL B cells were FACS sorted into 96-well PCR plates containing 2µl lysis mix per well. The index-sorting mode was activated to record the different fluorescence intensities of each sorted cell. Immediately after cell sorting, plates containing single cells in lysis mix were frozen on dry ice and stored at -80°C until further processing. For each plate, library preparation consisted in RT with template switching for incorporating well-specific barcodes and UMIs, cDNA amplification with 22 cycles of PCR, pooling of 96 wells into one tube, and 5’-end RNA-seq library preparation using tagmentation-based modified Nextera XT DNA sample Preparation kit (Illumina). Libraries were tagged with a plate-specific i7 index and were pooled by batches of 4-9 plates for sequencing on an Illumina NextSeq550 platform, with High Output 75-cycle flow cells, targeting 5×10^5^ reads per cell in paired-end single-index mode (Read 1: 67 cycles, Read i7: 8 cycles, Read 2: 16 cycles).

### 10x 3’ scRNA-seq

For experiments described in **Fig.1E**, frozen FL cell suspensions were thawed at 37°C in RPMI + 10% FCS, then washed and resuspended in FACS buffer (PBS + 5% FCS + 2 mM EDTA) at a 10^8^ cells/ml concentration. When cells from different donors were processed together in a same experiment (FL277, FL300, FL989, FL636) we performed cell hashing as described^71^. Cells were independently stained with a distinct hashtag oligonucleotide (HTO) conjugated antibody (Biolegend) for 30 min on ice (**Table S5**). Cells were then washed in PBS and independently stained with Live/Dead Fixable Aqua Dead Cell Stain (Thermofisher) for 10 min on ice. For each donor, total live cells were bulk-sorted by FACS with BD FACS Influx. Samples were pooled at equal concentration before encapsulation in 10x 3’ scRNA-seq. In some experiments (FL2, FL5, FL068, FL073, FL648, Tons104, Tons239), sorted human cells were pooled with mouse cell samples prior encapsulation in 10x 3’ scRNA-seq as described^36^.

Libraries were prepared with 10x Genomics Single Cell 3’ v2 workflow according to the manufacturer’s instructions. In experiments based on cell hashing, following cDNA amplification, SPRI select beads were used to separate the large cDNA fraction derived from cellular mRNAs (retained on beads) from the HTO-containing fraction (in supernatant). Each fraction was processed separately according to the 10x Genomics Single Cell 3’ v2 protocol to generate both transcriptome and HTO libraries. Resulting libraries were sequenced on an Illumina NextSeq550 platform with High Output 75-cycle flow cells, targeting 5×10^4^ reads per cell in paired-end single-index mode (Read 1: 26 cycles, Read i7: 8 cycles, Read 2: 57 cycles).

### 10x 5’ scRNA-seq

For experiments described in **Fig.1H**, frozen FL cell suspensions were thawed at 37°C in RPMI + 10% FCS, then washed and resuspended in FACS buffer (PBS + 5% FCS + 2 mM EDTA) at a concentration of 10^8^ cells/ml. Cells from different donors were processed together in two experiments (4 samples per experiment), using cell hashing as described^72^. Cells were independently stained with a distinct barcoded anti-human CD45 antibody (in-house conjugated) in FACS Buffer for 30 min on ice (**Table S5**). Cells were then washed in PBS and independently stained with Live/Dead Fixable Aqua Dead Cell Stain (Thermofisher) for 10 min on ice. For each donor, live cells were bulk-sorted by FACS with BD Influx. Samples were pooled at equal concentration, loaded for subsequent 10x Genomics Single Cell 5’ v1 workflow.

For the experiment described in **Fig.6D-E, S7, S8**, cultured cells were spun down and, in each well, cells were incubated with a mix containing a specific combination of barcoded anti-human CD45 antibodies (in-house conjugated, **Table S5, S6**) for 30 min on ice. Cells were washed in PBS, resuspended in FACS Buffer and all the wells were mixed together in a single tube. DAPI (Sigma-Aldrich) was added and live cells were sorted with the BD Influx cell sorter for subsequent 10x Genomics Single Cell 5’ v1 workflow.

### 10x 5’ scRNA-seq libraries were prepared according to the manufacturer’s instructions with modifications for generating the BCR-seq libraries

Following cDNA amplification, SPRI select beads were used to separate the large cDNA fraction derived from cellular mRNAs (retained on beads) from the HTO-containing fraction (in supernatant). For the cDNA fraction derived from mRNAs, 50ng were used to generate transcriptome library and around 10-20ng were used for BCR library construction. Gene expression libraries were prepared according to manufacturer’s instructions. For BCR libraries, heavy and light chain cDNA were amplified by two rounds of PCR (6 cycles + 8 cycles) using external primers recommended by 10x Genomics, and 800 pg of purified amplified cDNA was tagmented using Nextera XT DNA sample Preparation kit (Illumina) and amplified for 12 cycles using the SI-PCR forward primer (10x Genomics) and a Nextera i7 reverse primer (Illumina). For the HTO-containing fraction, 5ng were used to generate the HTO library.

The resulting libraries were pooled and sequenced together on an Illumina NextSeq550 platform, using High Output 75-cycle flow cells, targeting 5×10^4^ reads per cell for gene expression, 5×10^3^ reads per cell for BCR, 2×10^3^ reads per cell for hashtag, in paired-end single-index mode (Read 1: 26 cycles, Read i7: 8 cycles, Read 2: 57 cycles).

We also prepared and sequenced TCR-seq libraries from those experiments, but the resulting data were not analyzed in the results presented here.

### Immunohistochemistry staining

Immunohistochemistry protocol was followed to stain the FFPE tumor tissue samples of FL patients using antibodies listed in **Table S4**. Briefly, 3 µm sized paraffin embedded tissue sections were de-paraffinized with xylene then dehydrated through graded alcohols and subjected to antigen retrieval using HIER T-EDTA Buffer pH9 or pH6 (as indicated in **Table S4**) at 1X (Ref: ZUC029-500, Zytomed). Endogenous peroxidase activity was quenched with BLOXALL® Endogenous Blocking Solution (Ref: SP-6000, Vector) for 10 min in a humid room. Sections were washed with TBST (Tris Borate Saline Tween-20) and then blocked with Protein Block Serum-Free (Ref: No X0909, Dako) for 10 min. Slides were incubated with primary antibody diluted in Antibody Diluent (Ref: S2022, Dako) at concentrations indicated in **Table S4**. Slides were then washed for 3 min in TBST and incubated for 30 min with HRP (Horse Raddish Peroxidase) conjugated anti-rabbit (Ref: K4003, Dako) or anti-mouse (Ref: ENZ-ACC104-0150, Enzo) secondary antibody. After washing, slides were incubated with ImmPACT® DAB (Diaminobenzidine) (Ref: SK-4105, Vector). Color development were stopped in TBS 1X and washed in water. Slides were then counterstained with hematoxylin, mounted with VectaMount aqueous medium (Ref: H-5501, Vector) and observed under a light microscope (Olympus BX40).

Images were captured at 20X, using the Aperio Digital Pathology Slide Scanner, ScanScope XT of Leica. Images were viewed with Aperio ImageScope (Leica Biosystems).

### In vitro culture

Experiment was performed for FL338diag, FL338rel, FL068diag and FL068rel frozen FL cell suspensions. After cell staining with surface antibodies and Live/Dead Fixable Aqua Dead Cell Stain (Thermofisher), GC-like FL B cells were gated as CD3- CD14- IgD- CD38+ CD81+ single live cells and on the basis of λ or κ light chain isotype restriction. Mem-like FL B cells were gated as CD3- CD14- IgD- CD38- CD81- single live cells and on the basis of λ or κ light chain isotype restriction. Mid FL B cells were gated as CD3- CD14- IgD- CD38int CD81int single live cells and on the basis of λ or κ light chain isotype restriction. T_FH_ cells were gated as CD19- CD3+ CD4+ CD8- CD25- CXCR5+ PD1+ (**Fig.S7**). Bulk cell sorting was performed on BD FACS Influx.

For each donor, sorted GC-like, Mem-like and Mid FL B cells were cultured separately *in vitro* for 48 hours in a V-bottom 96-well plates in 200 µl complete RPMI medium at 37°C. Each well contained between 831 and 5,000 FL B cells depending on the cell number obtained after cell sorting. Cells were cultured alone, with sorted autologous T_FH_ cells (at a 1:1 ratio) or T_FH_ mimicking signals (including 0.02 μg/mL recombinant human IL-4, Peprotech; 0.05 μg/ml recombinant human IL-21, Peprotech, and 5 μg/mL anti-human CD40, Biolegend). Complete RPMI medium contained RPMI 1640 (Gibco) and was supplemented with 10% FCS, 1% glutamine, 100U/ml penicillin, 100 μg/ml streptomycin and 50 μM beta-mercaptoethanol. After 48 hours, cells were subjected to 10x 5’ scRNA-seq as described above.

### Spatial transcriptomics

The frozen FL tissue was retrieved from liquid nitrogen, shipped on dry ice, and stored at -80°C until processing. The tissue was embedded in OCT medium and a 10 µm section was cut on a cryostat and deposited on the capture area of the Visium slide following the guidelines of the 10x Genomics Visium Spatial Gene Expression protocol. The slide was stored at -80°C overnight, then processed for methanol fixation, hematoxylin and eosin staining, imaging, permeabilization (18 min), reverse transcription, second strand synthesis, denaturation, cDNA amplification (16 cycles of PCR), and library construction according to the manufacturer’s instructions (10x Genomics Visium Spatial Gene Expression). The resulting library was sequenced on an Illumina NextSeq2000, generating 170 million reads (Read 1: 28 cycles, Read i7: 10 cycles, Read i5: 10 cycles, Read 2: 79 cycles), yielding an average of 64,000 reads per spot for 2,647 tissue covered spots.

## COMPUTATIONAL METHODS

### Flow cytometry data analysis

#### FL B cells analysis

Frozen FL cell suspensions were thawed and stained following the same steps as described above in EXPERIMENTAL METHODS with the staining panel described in **Fig.S1C**. After acquisition, FCS files were imported into FlowJo software. Compensations were calculated by FACSDiva software using single-stained compensation tubes during acquisition and adjusted manually for each sample file.

Cells were gated on lymphocytes based on FSC and SSC values, doublets were eliminated and live cells were gated based on their LiveDead fluorescence intensity. For each sample, we performed a multi-parametric analysis. t-SNE plot was run on the live cell population using the t-SNE FlowJo plugin (based on default settings and all compensated parameters excluding LiveDead, FSC and SSC parameters). Gates were manually assigned to define non-malignant and malignant FL B cell clusters based on the specific fluorescence intensities of κ light chain, λ light chain, Dump (IgD, CD3, CD14), CD20 and CD10 surface markers. FL B cell clusters were identified based on their known isotype light chain restriction (**Table S2**), the high CD20 and CD10 expression levels and as being negative for the Dump. Non-malignant B cells were identified as B cells positive for either κ or λ light chain and as expressing lower CD20 and CD10 levels compared to their malignant counterparts. In the FL B cell population, we defined gates corresponding to GC-like, Mem-like or Mid cells based on CD38 expression values and on cut-offs defined on non-malignant B cells (**Fig.S1C**). For each population, we exported the Mean Fluorescence Intensity (MFI) for each marker used in the staining to analyze protein expression levels in each FL B cell subset. Relative MFI of every marker for each malignant subset was computed as the marker MFI of the malignant subset divided by the marker MFI of the non-malignant B cells for the same sample.

#### TME analysis

Frozen FL cell suspensions were thawed and stained following the same steps as described above in EXPERIMENTAL METHODS with the staining panel described in **Fig.S1D**. Samples were acquired on a BD LSR Fortessa. After acquisition, FCS files were imported into FlowJo software. Compensations were calculated by FACSDiva software using single-stained compensation tubes during acquisition and adjusted manually for each sample file. Cells were gated on lymphocytes based on FSC and SSC values, doublets were eliminated and live cells were gated based on their LiveDead fluorescence intensity. For each sample, a t-SNE plot was ran on the live cell population using the t-SNE FlowJo plugin (based on default settings and all compensated parameters excluding LiveDead, FSC, SSC parameters). Gates were manually assigned to define 6 cell subsets: **(1)** NK (Natural Killer), **(2)** activated and **(3)** unactivated CTL (cytotoxic T cells), **(4)** T_FH_ (T follicular helper cells), **(5)** T_REG_/T_FR_ (T regulatory/T follicular regulatory cells) and **(6)** T_N_/T_CM_ (naïve and central memory) CD4 T cells. Cell subsets were defined based on the specific fluorescence intensities of the markers used in the staining. **(1)** NK cells were defined as CD56+, CD3-, CXCR5-. **(2)** Activated CTL were defined as CD19- CD3+ CD8+ CD4-, PD1^hi^, CD127^lo^. **(3)** Unactivated CTL were defined as CD19- CD3+ CD8+ CD4-, PD1^lo^, CD127^hi^. **(4)** T_FH_ cells were defined as CD19- CD3+ CD4+ CD25- CXCR5+ PD1+. **(5)** T_REG_/T_FR_ cells were defined as CD19- CD3+ CD4+ CD25+ CD127^lo^. **(6)** T_N_/T_CM_ (naïve and central memory)

CD4 T cells were defined as CD19- CD3+ CD4+ excluding T_FH_ and T_REG_/T_FR_ cells. We then exported a CSV file containing for each donor cell numbers for each cell subsets. The percentage of each cell subpopulation among the live cell population was finally calculated for each donor.

### Pre-processing of scRNA-seq datasets

#### FB5P-seq

We used a custom bioinformatics pipeline to process fastq files and generate single-cell gene expression matrices and BCR sequence files as previously described^39^. Detailed instructions for running the FB5P-seq bioinformatics pipeline can be found at https://github.com/MilpiedLab/FB5P-seq. Quality control was performed on each dataset independently to remove poor quality cells. Cells with less than 250 genes detected were removed. We further excluded cells with values below 3 median absolute deviations (MADs) from the median for UMI counts, for the number of genes detected, or for ERCC accuracy, and cells with values above 3 MADs from the median for ERCC transcript percentage. For each cell, gene expression UMI count values were log-normalized with Seurat *NormalizeData* with a scale factor of 10,000^73^.

Index-sorting FCS files were visualized in FlowJo software and compensated parameters values were exported in CSV tables for further processing. For visualization on linear scales in the R programming software, we applied the hyperbolic arcsine transformation on fluorescence parameters^74^.

For BCR sequence reconstruction, the FB5P-seq pipeline used Trinity for *de novo* transcriptome assembly for each cell based on Read1 sequences, then MigMap for filtering the resulting contigs for productive BCR sequences and identifying germline V, D and J genes and CDR3 sequence for each contig. Filtered contigs were aligned to reference constant region sequences using Blastn. The FB5P-seq pipeline also ran the pseudoaligner Kallisto to map each cell’s FB5P-seq Read1 sequences on its reconstructed contigs and quantify contig expression. The outputs of the FB5P-seq pipeline were further processed and filtered with custom R scripts. For each cell, reconstructed contigs corresponding to the same V(D)J rearrangement were merged, keeping the largest sequence for further analysis. We discarded contigs with no constant region identified in-frame with the V(D)J rearrangement. In cases where several contigs corresponding to the same BCR chain had passed the above filters, we retained the contig with the highest expression level. BCR metadata from the MigMap and Blastn annotations were appended to the gene expression and index sorting metadata for each cell. Finally, heavy and light chain contig sequences were trimmed to retain only sequences from FR1 to the first 36 nucleotides of FR4 regions, and were exported as fasta files for further analysis of clonotypes and BCR phylogenies.

#### 10x 3’

Raw fastq files were processed using Cell Ranger software (v1.3.0 and v3.0.1), which performs alignment, filtering, barcode counting and unique molecular identifier (UMI) counting. In experiments where mixes of mouse and human cells were captured, reads were first aligned to the mm10 and GRCh38 genomes at the same time using Cell Ranger software, which resulted in barcodes assigned as murine or human or both. To improve alignment quality and avoid read loss due to mapping to both species, we then aligned cells only to the human GRCh38 reference, and filtered out the cell barcodes identified as mouse or human–mouse doublets in the dual alignment. In the experiment with hashtags, reads were directly aligned to the GRCh38 reference genome with CellRanger.

For each experiment, we retained cell barcodes if they had at least 500 genes with non-zero expression and genes if they were detected in at least 3 cells. We further excluded bad quality cells with values below 2 median absolute deviations (MADs) from the median for UMI counts or for the number of genes detected, and cells with values above 1.5 MADs for mitochondrial transcript percentage for most datasets. The threshold for percentage of mitochondrial transcripts was adjusted to 2 MADs for the dataset including hashtags. HTO barcodes for sample demultiplexing after hashing were counted using CITE-seq-count and were normalized for each cell using a centered log ratio (CLR) transformation across cells implemented in the Seurat function *NormalizeData*. Cells were demultiplexed using *MULTIseqDemux* function^75^ and barcodes assigned as doublets or negative were excluded from further analysis. The resulting filtered UMI count matrices were log-normalized with Seurat *NormalizeData* with a scale factor of 10,000.

#### 10x 5’

Raw fastq files from gene expression libraries were processed using Cell Ranger software (v3.0.1), with alignment on the GRCh38 reference genome. For each experiment, we retained cell barcodes if they had at least 500 genes with non-zero expression and genes if they were detected in at least 3 cells. We further excluded bad quality cells with values below 2 median absolute deviations (MADs) from the median for UMI counts or for the number of genes detected, and cells with values above 1.5-2 MADs for mitochondrial transcript percentage. HTO barcodes for sample demultiplexing after hashing were counted using CITE-seq-count and were normalized for each cell using a centered log ratio (CLR) transformation across cells implemented in the Seurat function *NormalizeData*. For experiments where samples were tagged with a single hashtag barcode, cells were demultiplexed using *MULTIseqDemux* function and barcodes assigned as doublets or negative were excluded from further analysis. For the experiment where samples were tagged with combinations of 3 hashtags (**Table S5** and **Fig.6D-E**), we manually set thresholds for hashtag positivity based on visual inspection of RidgePlots for all normalized hashtag signals, and selected cells which were positive for triplet combinations of hashtags used to label the different culture conditions. All other cells were excluded from further analysis. The resulting filtered UMI count matrices were log-normalized with Seurat *NormalizeData* with a scale factor of 10,000.

BCR-seq raw fastq files were processed with the FB5P-seq pipeline^39^ as described above for FB5P-seq datasets, omitting the part of the pipeline related to gene expression analysis, and using the list of cell-associated 10x barcodes from CellRanger analysis as inputs for splitting bam files upstream Trinity assembly of BCR contigs. BCR metadata from the MigMap and Blastn annotations were appended to the gene expression metadata for each cell. Heavy and light chain variable sequences, trimmed to retain only sequences from FR1 to the first 36 nucleotides of FR4 regions, were exported as fasta files for further analysis of clonotypes and BCR phylogenies.

### Segregation of TME, malignant and non-malignant B cells

For each sample in the 10x 3’ and 10x 5’ datasets, B cells were distinguished from TME non-B cells by applying low resolution graph-based cell clustering (Seurat *FindNeighbors* and *FindClusters*, resolution 0.1) and scoring B cell marker genes (*CD19, CD79A, CD79B, MS4A1*) with Seurat *AddModuleScore*. Clusters with a high B cell expression score were defined as B cells, other clusters were defined as TME non-B cells. In B cell clusters, non-malignant and malignant B cells were then distinguished by light chain isotype restriction. Briefly, cells were assigned as IGKC if the normalized expression of *IGKC* was superior to the sum of the normalized expressions of *IGLC* genes (*IGLC[2-7]*), or IGLC if the summed expression of *IGLC* genes was superior to the expression of *IGKC*. B cell clusters with balanced proportions of IGKC and IGLC cells were assigned as non-malignant, and clusters with IGKC or IGLC restriction were assigned as malignant. In 10x 5’ datasets, malignant B cell assignment was manually checked after identification of cells carrying the monoclonal malignant BCR rearrangement identified from single-cell BCR-seq analysis.

### Low dimension embedding

Variable genes (n=4000) were identified with Seurat *FindVariableFeatures (vst* method), BCR coding genes were excluded from the lists of variable genes. After centering with Seurat *ScaleData*, principal component analysis was performed on variable genes with Seurat *RunPCA*, and embedded in two-dimensional UMAP plots with Seurat *RunUMAP* on 40 principal components. UMAP embeddings colored by sample metadata were generated by Seurat *DimPlot*, those colored by single gene expression or module scores were generated by Seurat *FeaturePlot*, those colored by BCR sequence metadata or Score_GCtoMem_ and Score_TFH-activated_ were generated with ggplot2 *ggplot*.

### Computation of signature scores

We first clustered normal B cells analyzed by FB5P-seq on the basis of gene expression, discriminating GC, Mem and PC cells expressing expected marker genes (**Fig.S3A-C**). Clustering of the same cells on the basis of surface protein expression also segregated GC, Mem and PC cells (**Fig.S3D**). We retained cells characterized by the same identity in gene-based and protein-based clustering (**Fig.S3E**). To avoid overrepresentation of cell cycle-related genes in GC-specific signature, only non-cycling (G1 phase) GC B cells were used in those analyses (**Fig.S3F**), after cell cycle phases were attributed with Seurat *CellCycleScoring* using S or G2/M phases specific gene lists as previously described^76^. We used the *FindMarkers* function from Seurat package (test.use = “bimod”, min.diff.pct = 0.1, retaining only genes with adjusted p-value < 0.05) to detect differentially upregulated genes between **(1)** PC cells and non-PC cells (GC and Mem) (n = 710 genes), **(2)** non-PC cells and PC cells (n = 1313 genes), **(3)** GC cells and Mem cells (n = 690 genes), as well as **(4)** Mem cells and GC cells (n= 127 genes). The corresponding lists of genes are provided as **Supplementary Data Table 1**. For each one of the four signatures, signature scores were calculated for each single cell with Seurat *AddModuleScore*.

### Annotation of malignant B cells based on signature scores

Malignant B cell subsets were defined independently for FB5P-seq, 10x 3’, and 10x 5’ datasets. For each technology, cells passing QC (excluding cells from FL636, where too few malignant B cells were detected) were grouped into a single Seurat object, and expression scores corresponding to lists of signature genes were computed for every cell in the object using the Seurat function *AddModuleScore*. PC-like FL B cells were manually gated from malignant B cells with a threshold defined based on the separation of normal PC cells from non-PC cells (**Fig.2A-B**). After exclusion of PC-like cells, GC-like, Mid and Mem-like FL B cells were manually gated from malignant B cells with thresholds based on the separation of normal GC B cells from normal Mem B cells (**Fig.3A-B**).

For samples FL2 (FB5P-seq and 10x 3’ datasets) and FL539 (10x 5’ dataset), marker genes of PC-like or non-PC-like were computed with the Seurat *FindMarkers* (only.pos = TRUE, test.use = “bimod”) function on sample-specific malignant B cells. The resulting gene lists filtered by adjusted p-value < 0.05 are reported in **Supplementary Data Table 2**.

For all samples, marker genes of GC-like or Mem-like were computed with the Seurat *FindAllMarkers* (only.pos = TRUE, test.use = “bimod”, min.pct = 0.25, thresh.use = 0.25) function on sample-specific malignant non-PC-like B cells, and only marker genes with p < 0.05 were retained. Recurrent markers in the 21 samples analyzed by either FB5P-seq, 10x 3’ or 10x 5’ are reported in **Supplementary Data Table 3**.

Score_GCtoMem_ was computed independently for FB5P-seq, 10x 3’, and 10x 5’ datasets. For each dataset, a polynomial of degree 2 regression was fitted on the distribution of malignant B cells in the scatter plot of Mem (x-axis) and GC (y-axis) signature scores (**Fig.3B**), and the projection of data points on the regression curve was used to order cells from the most GC-like (Score_GCtoMem_ = 0) to the most Mem-like (Score_GCtoMem_ = 1) states.

Heatmaps were generated with custom scripts using the *pheatmap* function. Genes for PC-like heatmaps (**Fig.2D**) were selected on the basis of known expression in normal human PC. Genes for GC-like to Mem-like heatmaps (**Fig.3D**) were selected among top differentially expressed genes between GC-like and Mem-like FL B cells and on the basis of known expression in normal GC and Mem cells.

### Gene ontology

Gene ontology analysis (**Fig.2E**) was performed on the list of differentially expressed genes in FL539 PC-like cells, after removing *IGH* and *IGK/L* genes (**Supplementary Data Table 2**) with *enrichGO* function of the clusterProfiler R package, assessing “biological process” gene ontologies.

### Unsupervised analysis of intra-sample heterogeneity of malignant B cells

For each sample from the 10x 3’ and 10x 5’ datasets, we created a new Seurat object from malignant non-PC-like B cells only. After normalization and centering, we selected the top 4000 variable genes with *FindVariableFeatures (vst* method) and computed principal component analysis with *RunPCA*, followed by UMAP embedding with *RunUMAP* on 40 principal components. UMAP embeddings colored by Score_GCtoMem_ values are shown in **Fig.4A**.

Single-cell pseudo-time trajectories were computed with the DDRTree algorithm from the Monocle2 package in R^77^. Expression profiles were reduced to three dimensions using the *reduceDimension* function in the DDRTree package in R^78^. Analysis was performed as previously described^36^.

For each sample, we computed Pearson’s correlation coefficient between the cell loadings on the first principal component PC1 and Score_GCtoMem_ values (**Fig.4B-C**), and Pearson’s correlation coefficient between DDRTree pseudotime values and Score_GCtoMem_ values (**Fig.4D-E**).

### BCR-seq based phylogenies

For each sample processed with FB5P-seq or 10x 5’ scRNA-seq, we aligned single-cell IGL and IGH sequences to reference V_L_ and J_L_ as well as V_H_, D_H_ and J_H_ sequences respectively using IMGT HighV-quest^79^. Sequences which were not related to the major FL clonal rearrangement were removed from further analyses. We then inferred the unmutated common ancestor (UCA) sequence by combining the IMGT-defined germline V_L_, J_L_ and V_H_, D_H_, J_H_ sequences with the observed V_L_-J_L_ and V_H_-D_H_-J_H_ junctional sequences from the least somatically mutated FL B cell in the malignant clonotype. For each sample, the single-cell BCR repertoire metrics (*IGHV* gene, *IGHJ* gene, *IG(K/L)V* gene, *IG(K/L)J* gene, number of sequences, *IGHC* gene, number of mutations, number of N-glycosylation sites) for all cells and for gene expression subsets (GC-like, Mid, Mem-like, PC-like) are reported in **Supplementary Data Table 4**.

For phylogenetic analyses, concatenated variable (FR1 to FR4) IGH and IGL sequences of UCA and all clonally related FL cells were trimmed to equal length and aligned with the GCtree software^80^. Transcriptomic cell state (PC-like, GC-like, Mid, Mem-like) or sample of origin (FL338 diagnosis and FL338 relapse) were used as labels to color the different nodes and leaves of the resulting BCR sequence phylogenetic trees. For statistical analyses of label distribution among nodes and daughter leaves, we simulated random distributions of labels (cell state or sample of origin) in the same proportions as the observed distribution in the overall clonotype, and compared the proportion of labels between real and simulated trees for all large nodes in the tree (nodes with > 15 daughter leaves) with chi-squared tests. Adjusted p-values <0.05 are reported as * besides the corresponding node. The code used for statistical analyses of label distribution in GCtree BCR phylogenies is available upon request.

### TME analysis

For all samples in the 10x 3’ and 10x 5’ datasets, excluding FL636 (where too few malignant B cells were detected), we selected non-B TME cells passing quality controls, and constructed Seurat objects. Each Seurat object was log-normalized with *NormalizeData*, variable genes (n=4000) were identified with Seurat *FindVariableFeatures (vst* method), and BCR coding genes were excluded from the lists of variable genes. We integrated the sample-specific objects using the Seurat v3 method as described^73^, with *FindIntegrationAnchors* (anchor.features = 2000, dims = 1:40, k.filter = 92) and *IntegrateData* (dims = 1:40). The resulting integrated object was centered, reduced with *RunPCA* and embedded in two dimensions with *RunUMAP* on 40 principal components. The quality of the integration was assessed visually by *DimPlot* colored by sample origin, where cells from distinct samples were grouped in cell type-specific clusters.

After identifying cells in G1, S or G2/M phase of the cell cycle with *CellCycleScoring*, non-supervised graph-based clustering with *FindNeighbors* (k.param = 30) and *FindClusters* (resolution = 1) detected 15 clusters of TME cells (**Fig.6A**). We discarded clusters corresponding to doublets with B cells, unknown T cells, and S/G2/M cell cycle cells for further analyses. According to marker gene expression, we annotated 11 distinct TME cell subpopulations: **(1)** Naïve T cells, **(2)** Central Memory T helper cells, **(3)** T_REG_ (T regulatory) cells (combining T_REG_1 and T_REG_2 clusters), **(4)** T_FR_ (T follicular regulatory) cells, **(5)** T_FH_ (T follicular helper) cells type 1, **(6)** T_FH_ (T follicular helper) cells type 2, **(7)** Central T_FH_ cells, **(8)** Central Memory CTL (cytotoxic T) cells, **(9)** CTL effector cells, **(10)** T follicular CTL cells and **(11)** NK (Natural Killer) cells.

We then compared, for any given patient’s sample, the proportions obtained for each cell subset from FACS and scRNA-seq datasets. To achieve this, the 11 subpopulations identified by scRNA-seq were grouped in 6 subsets as follows: **(1)** Naïve T cells and Central Memory T helper cells were defined as CD4+ T_N_/T_CM_ cells, **(2)** T_REG_ and T_FR_ cells were defined as T_REG_/T_FR_ cells, **(3)** T_FH_ cells type 1, type 2, and Central T_FH_ cells were defined as T_FH_ cells, **(4)** Central Memory CTL were defined as CD8+ T_N_/T_CM_ cells, **(5)** CTL effector and T follicular CTL cells were defined as CD8+ CTL act. cells, and **(6)** NK cells were kept as originally identified. We then generated a CSV file containing, for each donor, cell numbers for each cell subset, and for all live cells. The percentage of each cell subpopulation among the live cell population was finally calculated for each donor. Linear regression and correlation analysis were performed between FACS and scRNA-seq cell proportions for each cell subset using Graphpad Prism.

### In vitro culture analysis

The demultiplexed 10×5’ dataset corresponding to the 4 samples cultured in 9 different conditions was loaded as a Seurat object. We identified non-B T_FH_ cells and malignant B cells as described above using marker genes signatures, light chain isotype restriction, and validation by BCR-seq information when available (FL338). Focusing only on malignant B cells, we annotated PC-like (n= 11 cells out of 4,315 total malignant B cells) and non-PC-like cells, and computed the Score_GCtoMem_ on non-PC-like cells as described above. We compared Score_GCtoMem_ values of cells initially sorted as different states (GC-like, Mid, Mem-like) and cultured in different conditions (FL B cells only, + auto-T_FH_, + aCD40 + IL4 + IL21), either for all samples pooled (**Fig.6E**) or for each sample individually (**Fig.S8A**), using Kruskal-Wallis test followed by pairwise Wilcoxon rank sum test with Benjamini-Hochberg p-value adjustment.

To identify genes that were robustly induced in malignant B cells by co-culture with auto-T_FH_, we computed for each sample (FL338 diag., FL338 rel., FL068 diag., FL068 rel.) and each initially sorted state (GC-like, Mid, Mem-like) the genes upregulated in the “+ auto-T_FH_” condition compared to the “FL B cells only” condition, using Seurat *FindMarkers* (only.pos = true, min.pct = 0.1, test.use = “bimod”), retaining only non-BCR coding genes with p-value < 0.01. For each initially sorted state, marker genes were ranked by recurrence in the 4 samples, and we retained genes with recurrence >1 as robust T_FH_-activated genes. The resulting 25-gene list (*PFN1, GRHPR, ISCU, PRDX2, UFC1, TXN, NFKBIA, CD40, CACYBP, GRN, CD83, PEA15, TESC, DHX9, TRAF4, PLIN3, APOBEC3G, NFKB2, CFLAR, ICAM1, CCDC28B, EBI3, MARCKS, CCL17, RGS1*) was used to compute the Score_TFH-activated_ with the Seurat function *AddModuleScore*. Comparison of Score_TFH-activated_ values in the “FL B cells only” and “+ auto-T_FH_” conditions allowed us to define the optimal threshold (Score_TFH-activated_ = 0.419, **Fig.S8B**) for identifying FL B cells having recently interacted with autologous T_FH_ cells, as shown in **Fig.6F**.

### Metaclustering

For each sample from the 10x 3’ and 10x 5’ datasets, we used the Seurat object of malignant non-PC-like B cells analyzed in the “unsupervised analysis of intra-sample heterogeneity of malignant B cells” as described above. Each sample was clustered with Seurat *FindNeighbors* (reduction = “pca”, dims = 1:40) and *FindClusters* (resolution = 0.8), and cluster-specific markers were identified with *FindAllMarkers* (test.use = “wilcox”, only.pos = true, min.pct = 0.1, logfc.threshold = 0.25) and filtered for p_val_adj < 0.05. Those analyses on 15 distinct datasets yielded 74 sample-specific graph-based malignant B cell clusters with 74 associated lists of marker genes. We then computed the similarity matrix of those 74 clusters by pairwise comparisons of lists of marker genes using the Jaccard similarity measure, computed as Jaccard(list1, list2) = length(intersect(list1, list2)) / length(union(list1, list2)). The resulting similarity matrix was reordered by hierarchical clustering using Rstats *hclust* (method = “ward.D2”), and the resulting dendrogram was used to define 6 metaclusters with *cutree* (k = 6), as shown in **Fig.6G**. Marker genes detected in each metacluster, ranked by the number of occurences they are found as markers of sample-specific cluster in a given metacluster, are reported in **Supplementary Data Table 5**.

Metaclusters were annotated as Mid, T_FH_-activated, T-B doublets, Mem-like, GC-like and Cell Cycle by inspection of the most recurrent marker genes (**Supplementary Data Table 5**) and by visualizing the distribution of Score_GCtoMem_, Score_CellCycle_ (computed with *AddModuleScore* on the list of genes used for defining S and G2/M cells), and Score_TFH-activated_ values on single cells grouped by metaclusters (**Fig.S8D-F**).

### Spatial Transcriptomics analysis

Spatial transcriptomics fastq files were first aligned on the GRCh38-3.0.0 reference by *SpaceRanger* v1.2.2 after manual registration of the H&E image using *Loupe browser* 5.0. The spot x gene matrix was loaded in the Seurat R package for subsetting the region of interest containing tumor follicles and normalization with *SCTransform*. We manually set the threshold to segregate nCount_SCT^hi^ follicular spots and nCount_SCT^lo^ non-follicular spots, and annotated spots by tissue area with the following rules: “centrofollicular” (CF) if a spot is follicular and all neighbor spots are follicular, “perifollicular” (PF) if a spot is follicular and at least one neighbor spot is non-follicular, “interfollicular” (IF) if a spot is non-follicular and at least one neighbor spot is follicular, “extrafollicular” (EF) if a spot is non-follicular and all neighbor spots are non-follicular, and “edge” if a spot is on the edge of the tissue section.

For deconvolution of cell type composition, we used the Python package *cell2location*^49^. We first constructed a reference scRNA-seq dataset by integrating all 10x 3’ and 10x 5’ datasets produced in the study (all cells passing quality controls, n = 15 samples) with the Seurat v3 method^73^ with *FindIntegrationAnchors* (default settings) and *IntegrateData* (default settings). The resulting integrated object was centered, reduced with *RunPCA* and embedded in two dimensions with *RunUMAP* on 30 principal components. We then used graph-based clustering with *FindNeighbors* (default settings) and *FindClusters* (resolution = 1.4), together with cell type or cell state annotations from our focused TME and metaclustering analyses, to annotate the integrated scRNA-seq reference with cell types as described in **Fig.7C**. Based on that annotation, lists of cell type-specific genes were computed with *FindAllMarkers* (test.use = “wilcox”, only.pos = TRUE, min.pct = 0.1, logfc.threshold = 0.25) and filtered for adjusted p-value < 0.01. We used those lists for computing signature scores in the scRNA-seq and spatial transcriptomics datasets using *AddModuleScore* as shown in **Fig.S9B** as an independent assessment of the *cell2location* output. For *cell2location* deconvolution, we reutilized the Jupyter notebooks provided by the authors for human lymph node analysis. That analysis outputs, for each spatial transcriptomics spot, a series of q05_spot_factors for every cell type annotated in the reference scRNA-seq dataset, corresponding to the probabilistic inference of the absolute number of cells of that cell type in that spot.

We then used the *cell2location* output spot x q05_spot_factors matrix in downstream visualization by *SpatialFeaturePlot*, manual gating and further quantification of spatial transcriptomics spots as described in **Fig.7D-G** and **Fig.S9B-C**. Differentially expressed genes in manually gated spot gates (**Fig.7E**) were computed with *FindAllMarkers* (test.use = “wilcox”, only.pos = TRUE, min.pct = 0.1, logfc.threshold = 0.25) and filtered for adjusted p-value < 0.05, then selected differentially expressed genes were visualized on a heatmap (**Fig.7H**) with pheatmap (scale = “row”).

### Statistical analysis

Statistical analyses were performed using Graphpad Prism or R softwares with tests and p-value significance criteria detailed in the figure legends.

## ACKNOWLEDGEMENTS

We are grateful to all past and present members of the Milpied and Nadel/Roulland laboratories at Centre d’Immunologie de Marseille-Luminy for useful discussions. We thank the Bioinformatics Core Facility of Centre d’Immunologie de Marseille-Luminy for helpful discussions and comments. We thank the Flow Cytometry Core Facility of Centre d’Immunologie de Marseille-Luminy. We are grateful to patients who consented their samples to the CeVi biobank, to pathologists from Institut Carnot CALYM CeVi centers, and to E. Mollaret from Institut Carnot CALYM for organizing samples transfer from the CeVi biobank. We acknowledge HalioDX and the UCA Genomix platform for sequencing. We acknowledge Centre de Calcul Intensif d’Aix-Marseille for granting access to its high performance computing resources. This work was supported by grants from Fondation ARC, Cancéropôle Provence-Alpes-Côte d’Azur, ANR (ANR-17-CE15-0009-01 JCJC MoDEx-GC), Institut Carnot CALYM and Institut Roche to P.M. This work was supported by institutional grants from INSERM, CNRS and Aix-Marseille University to the CIML. N.A. was supported by a fellowship from the French Ministry of Research and Higher Education.

## AUTHORSHIP CONTRIBUTIONS

N.A. designed experiments, performed experiments, analyzed the data, prepared the figures and wrote the manuscript. C.D. performed most bioinformatics analyses and prepared the figures. L.G. performed cell sorting, single-cell genomics and spatial transcriptomics experiments. I.C.M. performed initial bioinformatics analyses. T.G. performed bioinformatics analyses of the microenvironment. J.M.N. prepared key reagents. D.L.M. performed immunohistochemistry experiments. L.C. performed spatial transcriptomics experiment. F.L. supervised and helped with the analysis and interpretation of the immunohistochemistry experiments. P.G. collected FFPE samples, supervised and helped with the analysis and interpretation of the immunohistochemistry experiments. S.R. provided critical insight into data analysis and interpretation. L.S. supervised the development of bioinformatics and statistics methods. B.N. acquired funding and provided critical insight into data analysis and interpretation. P.M. designed experiments, performed experiments, analyzed the data, prepared the figures, wrote the manuscript, acquired funding and supervised the study. All authors revised the manuscript.

