## Supplementary Tables and Figures for "Functional plasticity and recurrent cell states of malignant B cells in follicular lymphoma"

| Sample origin | Sample name | Age | Sex | Datasets |
|---|---|---|---|---|
| Non-malignant spleen (deceased organ donor) | Sp18 | 52 | F | FB5P-seq |
| | Sp38 | na | na | FB5P-seq |
| | Sp29 | na | na | FB5P-seq |
| | Sp43 | na | na | FB5P-seq |
| Non-malignant tonsil (non-malignant reactive hyperplasia) | Tons061 | 30 | F | FB5P-seq |
| | Tons260 | 35 | M | FB5P-seq<br>10x 5’ |
| | Tons104 | 22 | M | 10x 3’ |
| | Tons239 | 41 | F | 10x 3’ |

na: not available

**Supplementary Table 1.** Characteristics of human non-malignant samples

| Sample name | Light chain | Age | Sex | Grade | Datasets |
|---|---|---|---|---|---|
| FL2 | κ | 68 | F | na | FB5P-seq<br>10x 3’ |
| FL5 | λ | na | M | na | FB5P-seq<br>10x 3’ |
| FL338diag | λ | 53 | M | 1-2 (diagnosis) | FB5P-seq<br>10x 5’<br>FACS<br>10x 5’ *in vitro* culture |
| FL338rel | λ | 54 | M | 3a (relapse) | FB5P-seq<br>10x 5’<br>FACS<br>10x 5’ *in vitro* culture |
| FL068rel | negative | 59 | M | na (relapse) | FB5P-seq<br>10x 3’<br>10x 5’<br>FACS<br>10x 5’ *in vitro* culture<br>IHC |
| FL068diag | negative | 56 | M | na (diagnosis) | 10x 5’<br>FACS<br>10x 5’ *in vitro* culture |
| FL281 | κ | 70 | F | 1-2 (relapse) | 10x 5’<br>FACS<br>FB5P-seq subset<br>IHC |
| FL296 | κ | 44 | M | 1-2 (diagnosis) | 10x 5’<br>FACS<br>FB5P-seq subset |
| FL539 | λ | 60 | F | 3a (relapse) | 10x 5’<br>FACS<br>FB5P-seq subset |
| FL636 | na | 37 | F | 3 (diagnosis) | 10x 3’ (Fig.1 then excluded from study) |
| FL277 | κ | 62 | M | 3a (diagnosis) | 10x 3’<br>FACS |
| FL989 | κ | 42 | F | 3a (relapse) | 10x 3’<br>FACS |
| FL648 | na | 68 | M | 1-2 (relapse) | 10x 3’<br>IHC |
| FL073 | κ | 60 | F | 1-2 (diagnosis) | 10x 3’<br>FACS<br>IHC |
| FL300 | λ | 51 | M | 1-2 (diagnosis) | 10x 3’<br>FACS<br>IHC |
| FL233 | κ | 66 | F | 3b (diagnosis) | FACS<br>IHC |
| FL_Visium1 | na | 28 | M | 1-2 (diagnosis) | Spatial Transcriptomics |

na: not available

**Supplementary Table 2.** Characteristics of human follicular lymphoma samples

| Antibody target | Fluorochrome | Clone | Manufacturer | Final dilution |
|---|---|---|---|---|
| BCL6 | PE CF594 | K112-91 | BD | 1/20 |
| CD3 | FITC | SK7 | Biolegend | 1/200 |
| CD3 | APC | UCHT1 | Biolegend | 1/50 |
| CD3 | APC-Cy7 | SK7 | Biolegend | 1/40 |
| CD4 | APC-Cy7 | RPA-T4 | Biolegend | 1/20 |
| CD8 | PE | SK1 | Biolegend | 1/100 |
| CD10 | PE | HI10a | Biolegend | 1/20 |
| CD14 | FITC | HCD14 | Biolegend | 1/200 |
| CD14 | APC-Cy7 | HCD14 | Biolegend | 1/40 |
| CD14 | APC-Cy7 | HCD14 | Biolegend | 1/40 |
| CD19 | BV785 | HIB19 | Biolegend | 1/100 |
| CD20 | PE-Cy7 | 2H7 | Biolegend | 1/40 |
| CD25 | BV650 | BC96 | Biolegend | 1/10 |
| CD27 | BV421 | M-T271 | Biolegend | 1/20 |
| CD38 | BV785 | HIT2 | Biolegend | 1/20 |
| CD44 | APC | BJ18 | Biolegend | 1/20 |
| CD44 | AF700 | BJ18 | Biolegend | 1/20 |
| CD56 | FITC | HCD56 | Biolegend | 1/10 |
| CD81 | PE-Dz594 | 5A6 | Biolegend | 1/20 |
| CD83 | PE-Dazzle594 | HB15e | Biolegend | 1/50 |
| CD127 | BV421 | A019D5 | Biolegend | 1/30 |
| CXCR4 | APC | 12G5 | Biolegend | 1/5 |
| CXCR5 | Biotin | RF8B2 | BD | 1/40 |
| CXCR5 | PerCP-Cy5.5 | RF8B2 | BD | 1/40 |
| GPR183 | FITC | SA313E4 | Biolegend | 1/20 |
| IgD | APC-Cy7 | IA6-2 | Biolegend | 1/20 |
| IgD | FITC | na | Invitrogen | 1/100 |
| Ki67 | APC | KI-67 | Biolegend | 1/20 |
| κ light chain | BV421 | TB28-2 | Biolegend | 1/20 |
| κ light chain | PerCP-Cy5.5 | G20-193 | BD | 1/20 |
| λ light chain | BV421 | JDC-12 | BD | 1/20 |
| λ light chain | PerCP-Cy5.5 | MHL-38 | Biolegend | 1/20 |
| PD1 | PE-Cy7 | NAT105 | Biolegend | 1/20 |
| Streptavidin | PE | | Biolegend | 1/2000 |

**Supplementary Table 3.** Antibodies used for flow cytometry and cell sorting

| Target | Clone | Species | Reference, Manufacturer | Antigen Unmasking | Dilution |
|---|---|---|---|---|---|
| CD69 | EPR21814 | rabbit | ab233396, Abcam | pH9 | 1/500 |
| CD81 | polyclonal | rabbit | HPA007234, Merck | pH9 | 1/200 |
| BCL6 | LN22 | mouse | PA0204, Leica | pH9 | RTU |
| NANS | B-6 | mouse | sc-374133, Santa Cruz | pH6 | 1/200 |
| CD38 | SPC32 | mouse | NCL-L-CD38-290, Leica | pH6 | 1/100 |
| Ki67 | 30-9 | rabbit | 790-4286, Ventana | pH9 | RTU |
| CD3 | F7.2.38 | mouse | M725401, Dako | pH9 | 1/50 |
| CD5 | 4C7 | mouse | NCL-L-CD5-4C7, Leica | pH9 | 1/100 |
| CD21 | 2G9 | mouse | NCL-L-CD21-2G9, Leica | pH6 | 1/100 |

**Supplementary Table 4.** Antibodies and conditions used for immunohistochemistry staining. RTU: ready to use.

| Hashtag antibody | Target | Clone | Manufacturer | Barcode sequence (5’-3’) | Final dilution |
|---|---|---|---|---|---|
| TotalSeqA-0251 | CD298 / β2M | LNH-94 / 2M2 | Biolegend | GTCAACTCTTTAGCG | 1/500 |
| TotalSeqA-0252 | CD298 / β2M | LNH-94 / 2M2 | Biolegend | TGATGGCCTATTGGG | 1/500 |
| TotalSeqA-0253 | CD298 / β2M | LNH-94 / 2M2 | Biolegend | TTCCGCCTCTCTTTG | 1/500 |
| TotalSeqA-0254 | CD298 / β2M | LNH-94 / 2M2 | Biolegend | AGTAAGTTCAGCGTA | 1/500 |
| 5pHTO_BC1_PMv2 | CD45 | HI30 | Biolegend / in-house conjugated | TGCCATAGGTAC | 1/20 |
| 5pHTO_BC2_PMv2 | CD45 | HI30 | Biolegend / in-house conjugated | TGAAGAACGCCG | 1/20 |
| 5pHTO_BC3_PMv2 | CD45 | HI30 | Biolegend / in-house conjugated | CCGCACTGGAGT | 1/20 |
| 5pHTO_BC4_PMv2 | CD45 | HI30 | Biolegend / in-house conjugated | AAGGTCACAGCT | 1/20 |
| 5pHTO_BC5_PMv2 | CD45 | HI30 | Biolegend / in-house conjugated | CCATTCGCGATA | 1/20 |
| 5pHTO_BC6_PMv2 | CD45 | HI30 | Biolegend / in-house conjugated | GTCTCAACTGCG | 1/20 |
| 5pHTO_BC7_PMv2 | CD45 | HI30 | Biolegend / in-house conjugated | AGCCAATATTCT | 1/20 |
| 5pHTO_BC8_PMv2 | CD45 | HI30 | Biolegend / in-house conjugated | AGGACCGTTAGC | 1/20 |
| 5pHTO_BC9_PMv2 | CD45 | HI30 | Biolegend / in-house conjugated | AGCGTCTCGTTC | 1/20 |
| 5pHTO_BC10_PMv2 | CD45 | HI30 | Biolegend / in-house conjugated | TGGAGATCCACG | 1/20 |
| 5pHTO_BC11_PMv2 | CD45 | HI30 | Biolegend / in-house conjugated | ATGCGCTATGGC | 1/20 |
| 5pHTO_BC12_PMv2 | CD45 | HI30 | Biolegend / in-house conjugated | CACTTGGATTAA | 1/20 |

**Supplementary Table 5. “**Hashtag” antibodies used for sample multiplexing in 10x 3’ and 10x 5’ scRNA-seq experiments.

| Sample name | Culture Condition | anti-CD45 “hashtag” 1 | anti-CD45 “hashtag” 2 | anti-CD45 “hashtag” 3 |
|---|---|---|---|---|
| **FL068 diag** | GC-like only | BC4 | BC7 | BC11 |
| | Mem-like only | BC5 | BC7 | BC11 |
| | Mid only | BC6 | BC7 | BC11 |
| | GC-like + auto-$T_{FH}$ | BC1 | BC7 | BC11 |
| | Mem-like + auto-$T_{FH}$ | BC2 | BC7 | BC11 |
| | Mid + auto-$T_{FH}$ | BC3 | BC7 | BC11 |
| | GC-like only | BC3 | BC7 | BC12 |
| | Mem-like only | BC1 | BC7 | BC12 |
| | Mid only | BC2 | BC7 | BC12 |
| | GC-like + $T_{FH}$-like | BC6 | BC7 | BC12 |
| | Mem-like + $T_{FH}$-like | BC4 | BC7 | BC12 |
| | Mid + $T_{FH}$-like | BC5 | BC7 | BC12 |
| **FL068 rel** | GC-like only | BC1 | BC8 | BC11 |
| | Mem-like only | BC2 | BC8 | BC11 |
| | Mid only | BC3 | BC8 | BC11 |
| | GC-like + auto-$T_{FH}$ | BC4 | BC8 | BC11 |
| | Mem-like + auto-$T_{FH}$ | BC5 | BC8 | BC11 |
| | Mid + auto-$T_{FH}$ | BC6 | BC8 | BC11 |
| | GC-like only | BC3 | BC8 | BC12 |
| | Mem-like only | BC1 | BC8 | BC12 |
| | Mid only | BC2 | BC8 | BC12 |
| | GC-like + $T_{FH}$-like | BC6 | BC8 | BC12 |
| | Mem-like + $T_{FH}$-like | BC4 | BC8 | BC12 |
| | Mid + $T_{FH}$-like | BC5 | BC8 | BC12 |
| **FL338 diag** | GC-like only | BC1 | BC9 | BC11 |
| | Mem-like only | BC2 | BC9 | BC11 |
| | Mid only | BC3 | BC9 | BC11 |
| | GC-like + auto-$T_{FH}$ | BC4 | BC9 | BC11 |
| | Mem-like + auto-$T_{FH}$ | BC5 | BC9 | BC11 |
| | Mid + auto-$T_{FH}$ | BC6 | BC9 | BC11 |
| | GC-like only | BC3 | BC9 | BC12 |
| | Mem-like only | BC1 | BC9 | BC12 |
| | Mid only | BC2 | BC9 | BC12 |
| | GC-like + $T_{FH}$-like | BC6 | BC9 | BC12 |
| | Mem-like + $T_{FH}$-like | BC4 | BC9 | BC12 |
| | Mid + $T_{FH}$-like | BC5 | BC9 | BC12 |

| | | | | |
|---|---|---|---|---|
| **FL338 rel** | GC-like only | BC4 | BC10 | BC11 |
| | Mem-like only | BC5 | BC10 | BC11 |
| | Mid only | BC3 | BC10 | BC11 |
| | GC-like + auto-$T_{FH}$ | BC1 | BC10 | BC11 |
| | Mem-like + auto-$T_{FH}$ | BC2 | BC10 | BC11 |
| | Mid + auto-$T_{FH}$ | BC6 | BC10 | BC11 |
| | GC-like only | BC3 | BC10 | BC12 |
| | Mem-like only | BC4 | BC10 | BC12 |
| | Mid only | BC5 | BC10 | BC12 |
| | GC-like + $T_{FH}$-like | BC6 | BC10 | BC12 |
| | Mem-like + $T_{FH}$-like | BC1 | BC10 | BC12 |
| | Mid + $T_{FH}$-like | BC2 | BC10 | BC12 |

**Supplementary Table 6.** Combinations of anti-CD45 “hashtag” antibody barcodes (5pHTO_BCx_PMv2) used for 10x 5’ scRNA-seq after *in vitro* culture.

| Sample name | scRNA-seq dataset | GC-like (%) | Mid (%) | Mem-like (%) | PC-like (%) |
|---|---|---|---|---|---|
| FL068rel | FB5P-seq | 21.15 | 64.4 | 14.25 | 0 |
| FL2 | | 0.52 | 80.93 | 15.46 | 3.09 |
| FL338diag | | 6.55 | 76.21 | 17.24 | 0 |
| FL338rel | | 7.78 | 90.42 | 1.8 | 0 |
| FL5 | | 29.5 | 70.5 | 0 | 0 |
| FL2 | 10x 3’ | 0.73 | 74.18 | 22.9 | 2.18 |
| FL277 | | 9.1 | 51.99 | 38.92 | 0 |
| FL300 | | 4.18 | 74.26 | 21.57 | 0 |
| FL5 | | 26.30 | 70.6 | 3.02 | 0.07 |
| FL989 | | 6.06 | 82.7 | 11.23 | 0 |
| FL068rel | | 21.78 | 55.34 | 22.82 | 0 |
| FL073 | | 7.68 | 82.87 | 9.45 | 0 |
| FL648 | | 16.07 | 81.83 | 1.84 | 0.26 |
| FL068diag | 10x 5’ | 16.44 | 71.11 | 12.44 | 0 |
| FL068rel | | 24.23 | 66.7 | 8.87 | 0.21 |
| FL338diag | | 7.13 | 66.57 | 26.3 | 0 |
| FL338rel | | 13.35 | 69.17 | 17.43 | 0.047 |
| FL281 | | 54.39 | 45.14 | 0.47 | 0 |
| FL296 | | 8.13 | 89.98 | 1.78 | 0.11 |
| FL539 | | 5.35 | 29.72 | 62.58 | 2.36 |

**Supplementary Table 7.** Proportions of malignant FL B cell subsets in FL samples

Data are presented as the percentage of GC-like, Mid, Mem-like or PC-like cells among malignant FL B cells for each sample, as defined by scRNA-seq analysis.

### Supplementary Figure 1

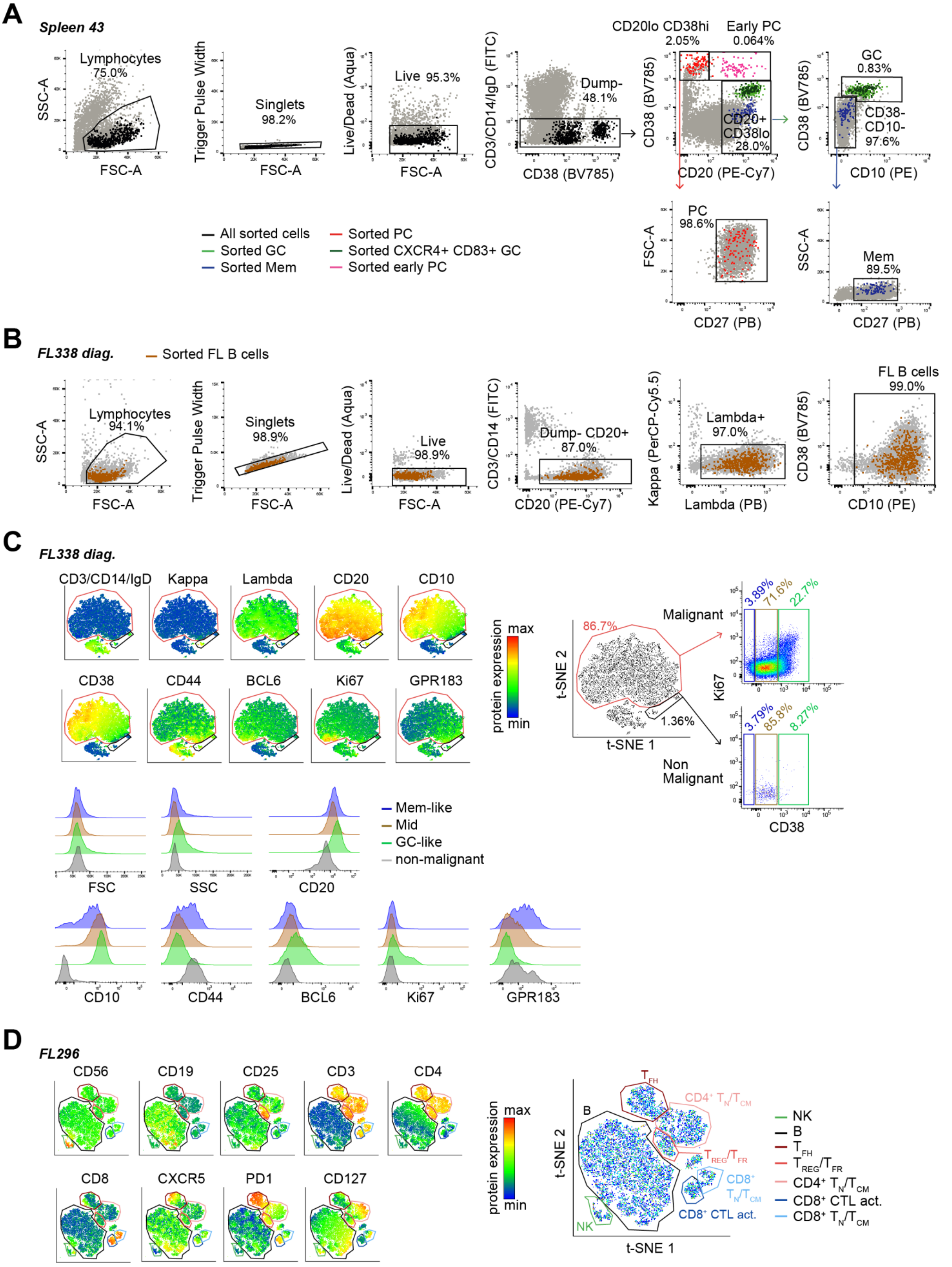

##### Supplementary Figure 1. FACS sorting and gating strategies.

**(A)** Related to **Fig.1A**. Flow cytometry gating strategy for normal single B cell sort from frozen human cell suspensions for spleen or tonsil samples for FB5P-seq, here on the example of Sp43. Sorted cells are labeled as colored dots over grey dots (total sample cells), using the indicated color code. Numbers above gates indicate the percentage of cells in the indicated gate relative to its parent gate. **(B)** Related to **Fig.1B**. Flow cytometry gating strategy for FL B cell single-cell sort from frozen FL cell suspensions for FB5P-seq, here on the example of FL338 diag. Sorted cells are labeled as brown dots over grey dots (total FL cells). Numbers above gates indicate the percentage of cells in the indicated gate relative to its parent gate in the total FL sample. **(C)** Related to **Fig.3F-G**. Flow cytometry gating strategy for the identification of malignant and non-malignant B cells in FL samples, and subsequent GC-like, Mid and Mem-like FL B cell states identification, here on the example of FL338 diag. A t-SNE embedding was computed from the expression levels of the indicated 10 fluorescent markers. Malignant and non-malignant B cell clusters were gated on the t-SNE plot. GC-like, Mid and Mem-like malignant B cells were gated based on CD38 levels. Fluorescence intensities were extracted for each marker in GC-like, Mid, Mem-like and non-malignant B cells for preparing **Fig.3G**. **(D)** Related to **Fig.6A-C**. Flow cytometry gating strategy for the identification of natural killer (NK) cells, B cells, T follicular helper ($T_{FH}$) cells, T regulatory/T follicular regulatory ($T_{REG}/T_{FR}$) cells, naïve and/or central memory $CD4^+$ T cells (CD4+ TN/TCM), naïve and/or central memory $CD8^+$ T cells (CD8+ TN/TCM), and activated cytotoxic $CD8^+$ T cells (CD8+ CTL act.) in FL cell suspensions, here on the example of FL296. A t-SNE embedding was computed from the expression levels of the indicated 9 fluorescent markers. TME subsets were gated on the t-SNE plot.

### Supplementary Figure 2

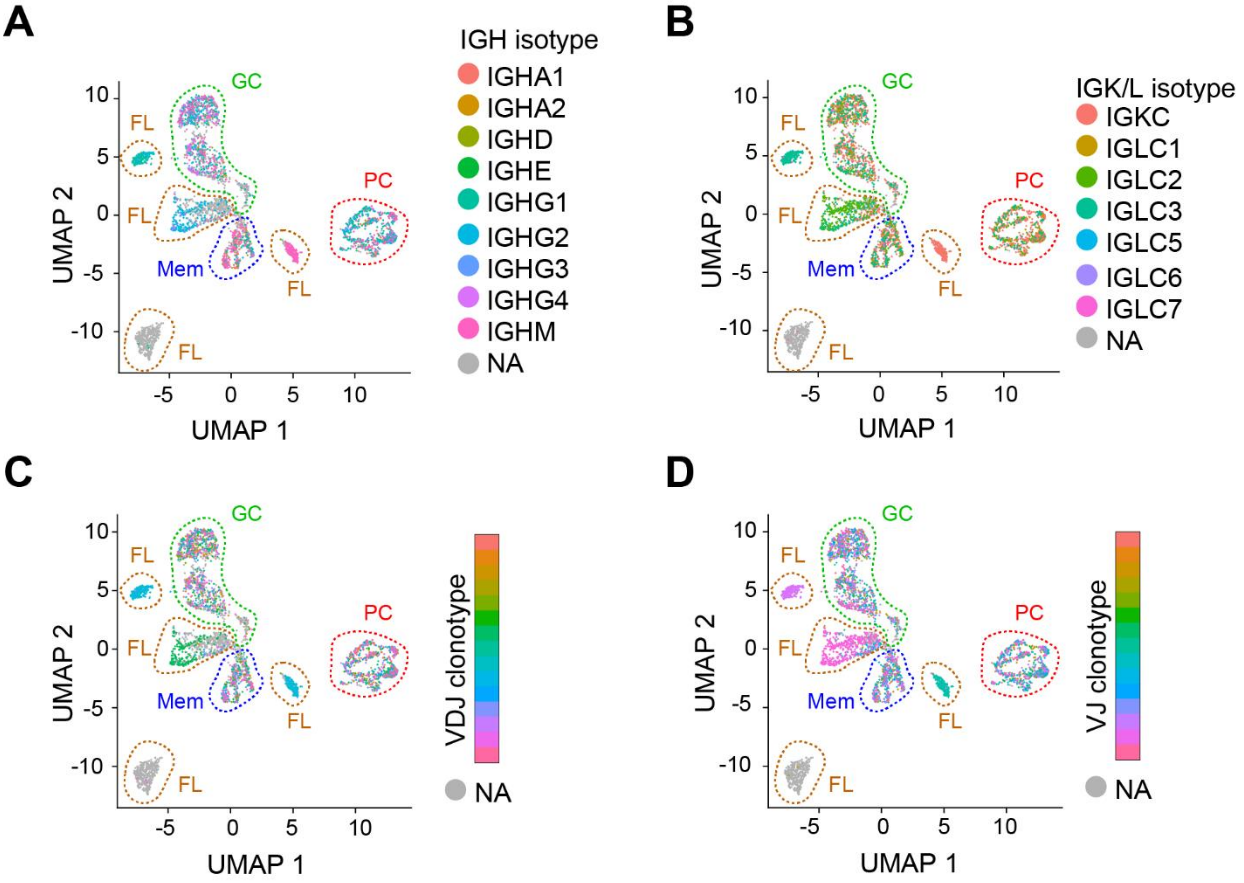


**Supplementary Figure 2. Isotype and BCR clonality of normal and FL B cells analyzed by FB5P-seq**

Related to **Fig.1C-D**. **(A,B)** UMAP embedding of normal and FL B cells analyzed by FB5P-seq. Cells are colored based on **(A)** heavy chain (IGH) or **(B)** light chain (IGK/L) isotype. **(C,D)** UMAP embedding of normal and FL B cells analyzed by FB5P-seq. Cells are colored based on **(C)** $V_H$-$D_H$-$J_H$ or **(D)** $V_L$-$J_L$ clonotypic combinations. *n*=3,722 B cells. NA= not applicable, i.e., no IGH or no IGK/L sequence reconstructed. Clusters of cells corresponding to the indicated cell types are circled by dashed colored lines.

#### Supplementary Figure 3

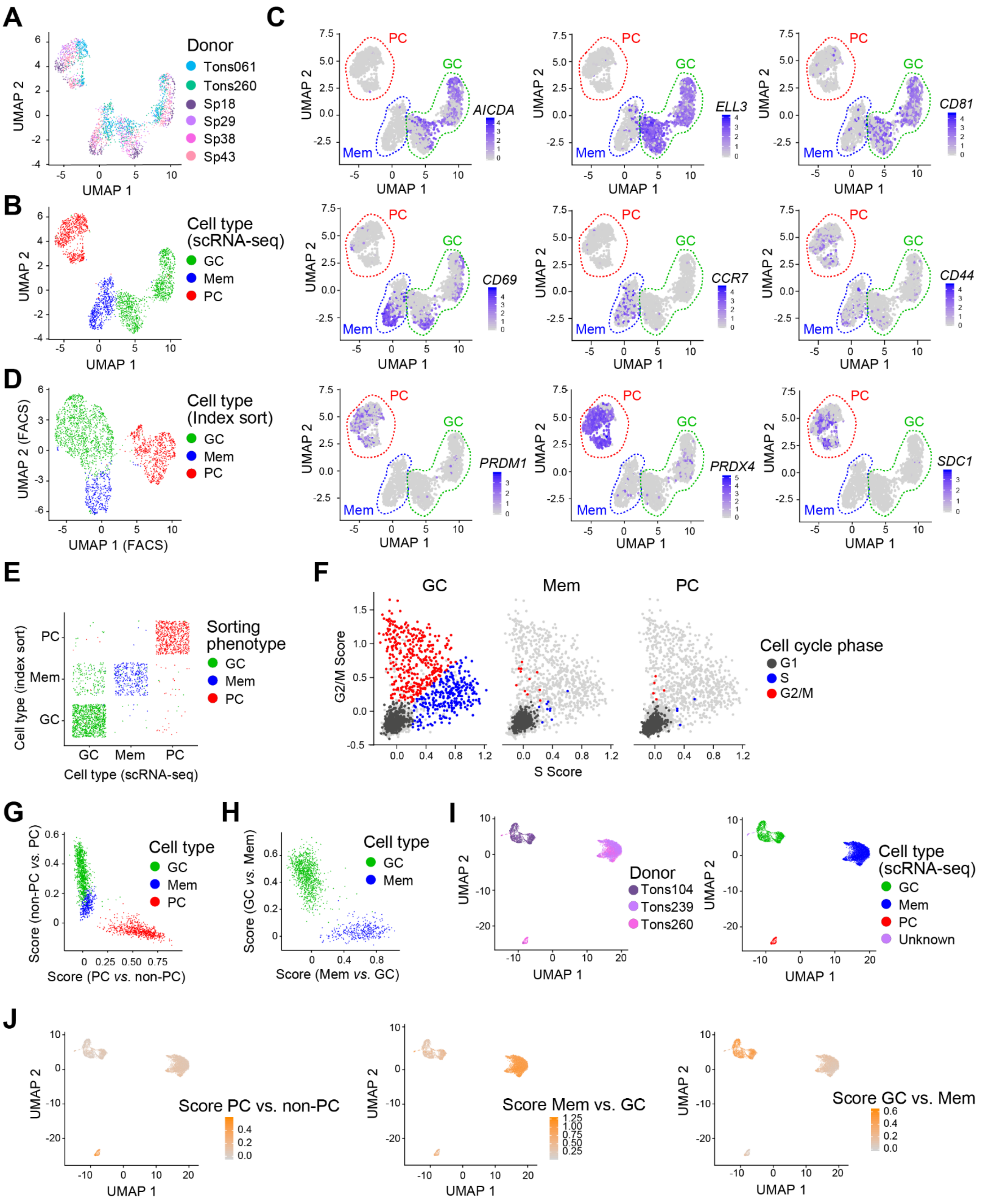


**Supplementary Figure 3. Construction and validation of normal B cell signatures**

Related to **Fig.2** and **Fig.3**. **(A-B)** UMAP embedding of single normal spleen and tonsil B cells analyzed by FB5P-seq. Cells are colored by donor origin **(A)** or cell type (defined by transcriptome analysis) **(B)**.

**(C)** Feature plots of normal B cells from the FB5P-seq dataset in UMAP embedding. Cells are colored based on the expression of GC (*AICDA*, *ELL3*, *CD81*, top row), Mem (*CD69*, *CCR7*, *CD44*, middle row), or PC (*PRDM1*, *PRDX4*, *SDC1*, bottom row) cell-specific marker genes as indicated. Clusters of cells corresponding to the indicated cell types are circled by dashed colored lines. **(D)** UMAP embedding of normal B cells from the FB5P-seq dataset, based only on index sorting surface protein parameters. Cells are colored according to cell type defined by the index sorting re-analysis. **(E)** Intersection of index sorting-defined and scRNA-seq-defined cell type identity for each normal B cell from the FB5P-seq dataset. Cells are colored based on their initial sorting phenotype. **(F)** Scatter plots of G2/M score *vs.* S score for normal GC, Mem or PC cells from the FB5P-seq dataset, identified based on the intersection between the scRNA-seq cell type and the index sort cell-type as presented in **(E)**. Only GC cells in G1 phase were considered for generating GC-specific gene signatures. **(G)** Non-PC *vs.* PC signature score plotted against PC *vs.* non-PC signature score for single normal spleen and tonsil B cells (FB5P-seq) colored by cell type, presented as a scatter plot. **(H)** GC *vs.* Mem signature score plotted against Mem *vs.* GC signature score for single normal spleen and tonsil GC and Mem B cells (FB5P-seq) colored by cell type, presented as a scatter plot. **(I-J)** UMAP embedding of tonsil B cells from the 10x 3' and 10x 5' scRNA-seq normal B cell datasets, after integration of the 3 datasets. Cells are colored by **(I)** donor (left) and cell type (right, defined by scRNA-seq analysis), or **(J)** PC vs. non-PC signature score (left), Mem *vs.* GC signature score (middle), or GC *vs.* Mem signature score (right).

#### Supplementary Figure 4

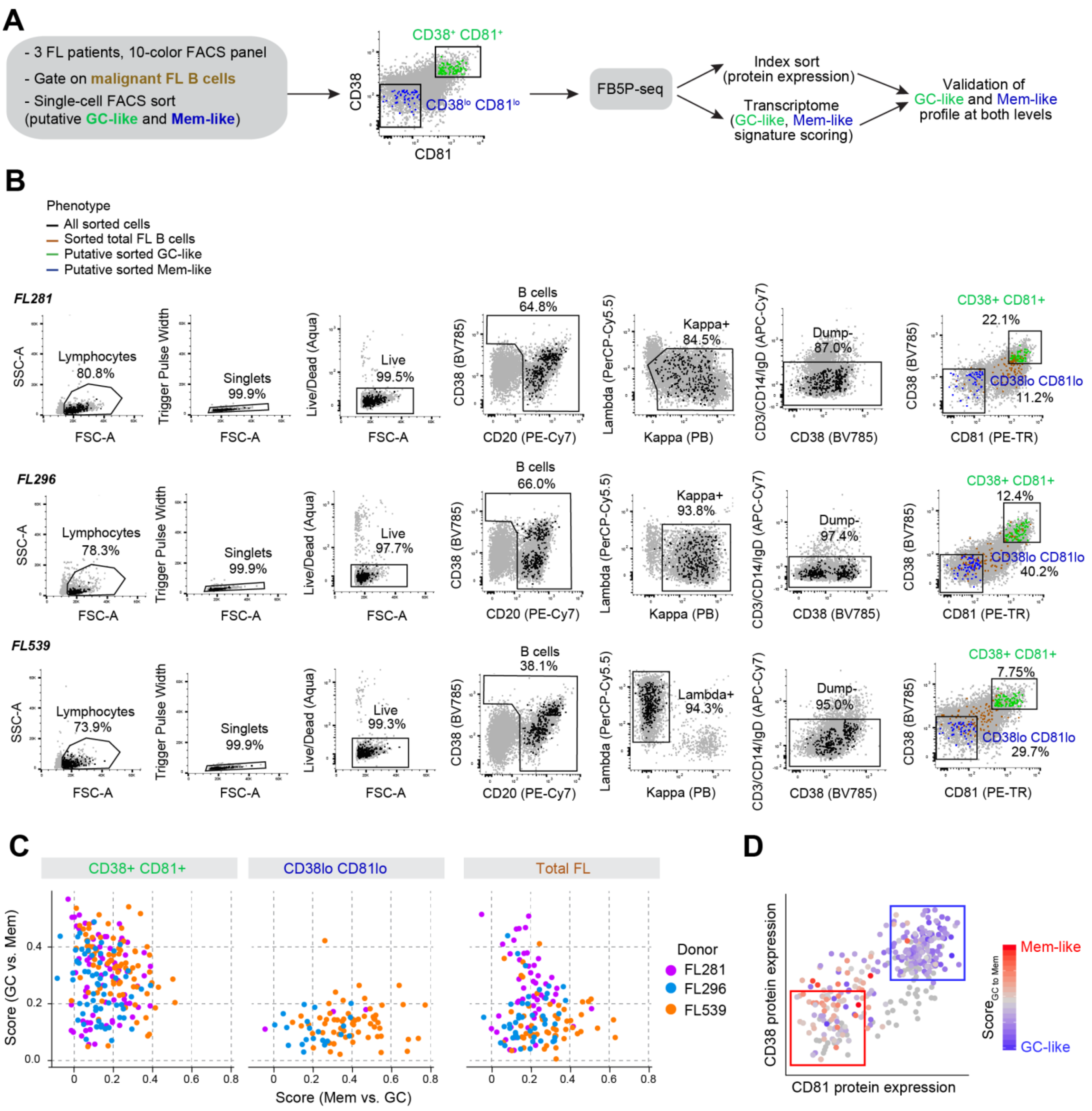


**Supplementary Figure 4. CD38$^{hi}$CD81$^{hi}$ and CD38$^{lo}$CD81$^{lo}$ phenotypes identify GC-like and Mem-like FL B cell states, respectively**

Related to **Fig.3F-G**. **(A)** Experimental workflow for putative GC-like and Mem-like FL B cell sort and subsequent FB5P-seq analysis. **(B)** Flow cytometry gating strategy for total FL B cells as well as putative GC-like and Mem-like single-cell sorting from FL cell suspensions for samples FL281, FL296 and FL539. Sorted cells are labeled as colored dots over grey dots (total sample cells), using the color code indicated at the top (Phenotype). Numbers above gates indicate the percentage of cells in the indicated gate relative to its parent gate. **(C)** Scatter plots of GC *vs.* Mem gene expression signature score plotted against Mem *vs.* GC gene expression signature score for cells sorted as putative GC-like (left), Mem-like (middle), or total (right) FL B cells from FL281, FL296 and FL539 samples, as indicated

(Donor). Each dot represents an individual cell. **(D)** Scatter plot of surface protein expression of CD38 *vs.* CD81 on all FL B cells. Each symbol represents an individual cell. Cells were colored according to their gene expression profile along the GC-like to Mem-like continuum ($Score_{GCtoMem}$).

#### Supplementary Figure 5

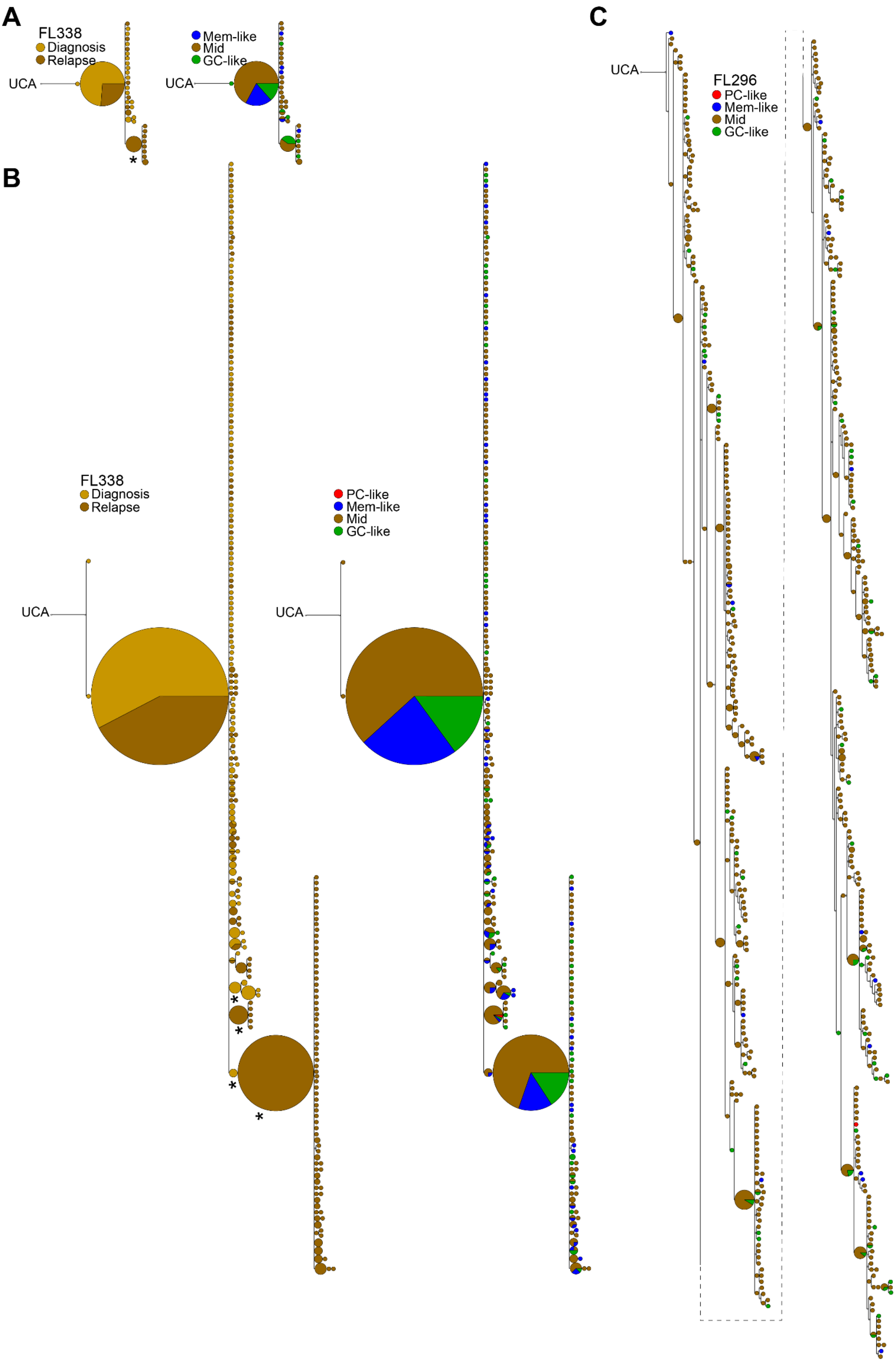

**Supplementary Figure 5. GC-like, Mem-like and PC-like FL B cell states are independent of subclonal divergence**

Related to **Fig.5**. **(A-C)** Phylogenetic trees built from concatenated *IGH-IGK/L* nucleotide sequences of clonally related single malignant FL B cells for samples FL338 (diag. and rel. samples are grouped) analyzed by FB5P-seq **(A)** or 10x 5' **(B)**, and sample FL296 **(C)**. The non-filled circle symbol (top left) indicates the inferred UCA sequence (unmutated common ancestor) to which the tree is rooted. Each symbol in the tree corresponds to a unique *IGH-IGK/L* sequence; symbol size is relative to the number of cells carrying that sequence. Symbols are colored according to donor origin (diag. or rel. in **(A-B)**) or cell subset (PC-like: red, GC-like: green, Mem-like: blue, Mid: brown) **(A-C)**. In the case of multiple cells carrying the same sequence, proportions of cells in each state are indicated as a pie chart within the symbol. * besides a node indicates a statistically significant difference in cell origin or states (colors) proportions among cells below that node compared to the whole sample.

### Supplementary Figure 6

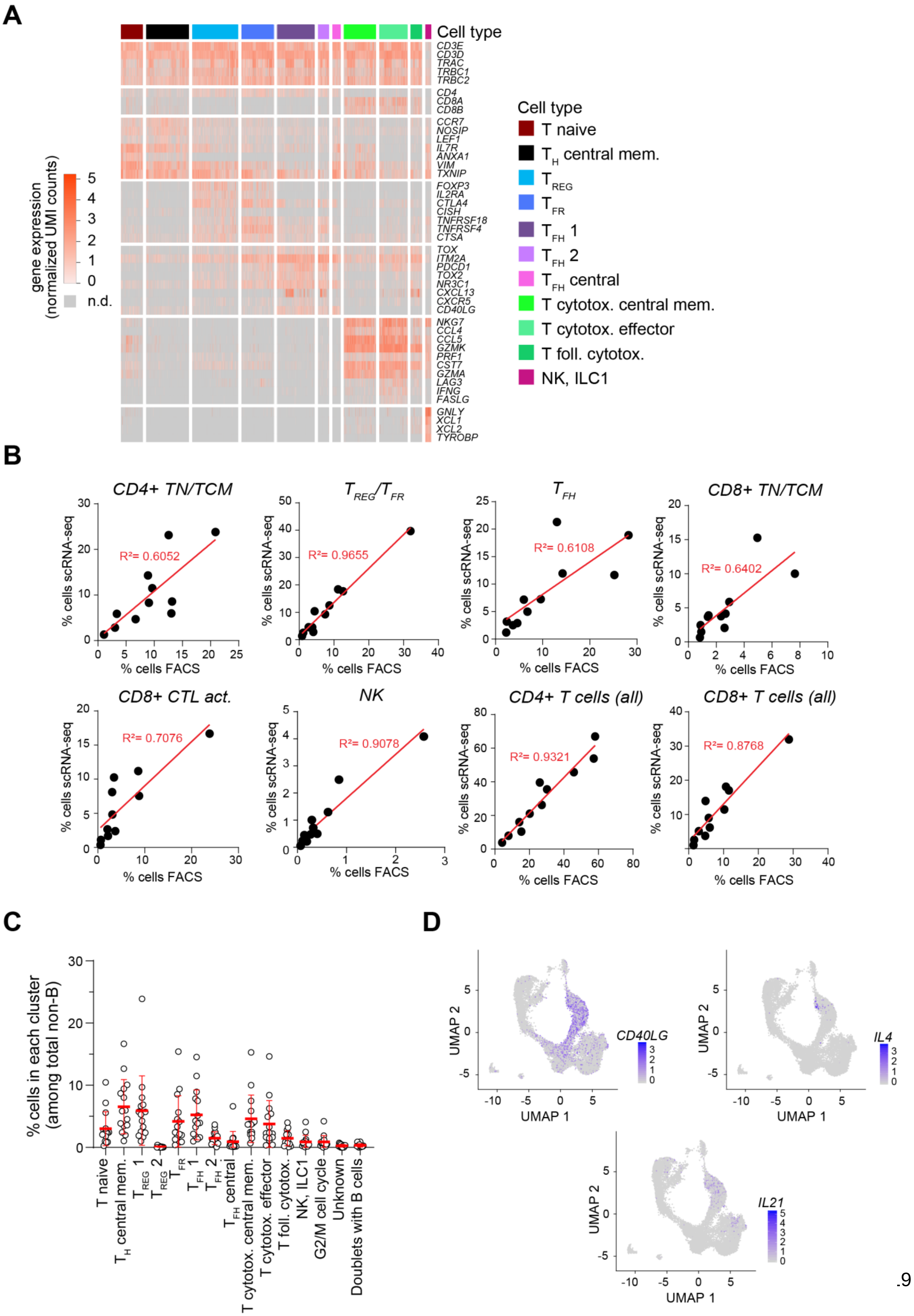

**Supplementary Figure 6. Quantification of FL TME cell types by flow cytometry and scRNA-seq**

Related to **Fig. 6A-C**. **(A)** Gene expression heatmap of single TME non-B cells for the expression of indicated marker genes used to annotate unsupervised clusters. Cells (columns) are grouped by cell type clusters as indicated by the bar above plot. Genes (rows) are grouped, from top to bottom, as pan T cell markers, *CD4* and *CD8* genes, naïve and central memory markers, $T_{REG}$ markers, $T_{FH}$ markers, CTL markers, and NK markers. **(B)** Scatter plots of cell type percentage among live cells as determined by flow cytometry (x-axis) or scRNA-seq (y-axis) in FL samples for the indicated cell types: CD4$^+$ naive/central memory T cells (CD4+ TN/TCM), T regulatory/T follicular regulatory cells ($T_{REG}$/$T_{FR}$), T follicular helper cells ($T_{FH}$), CD8$^+$ naive/central memory T cells (CD8+ TN/TCM), CD8$^+$ activated cytotoxic T cells (CD8+ CTL act.), natural killer cells (NK), total CD4+ T cells and total CD8+ T cells. $R^2$: goodness of fit coefficient for linear regression. Each symbol represents an individual sample ($n$=11 samples). **(C)** Proportions of cell subsets defined by graph-based clustering in scRNA-seq datasets (shown in **Fig.6A**), computed as the percentage of cells in each cluster among total TME non-B cells for each sample. Each symbol represents an individual sample ($n$=15 samples). **(D)** Feature plots of TME non-B cells UMAP embedding presented in **Fig.6A**. Cells are colored based on the expression of *CD40LG* (top left), *IL4* (top right), or *IL21* (bottom).

### Supplementary Figure 7

## A

**Follicular Lymphoma B cells sorting strategy for *in vitro* culture**

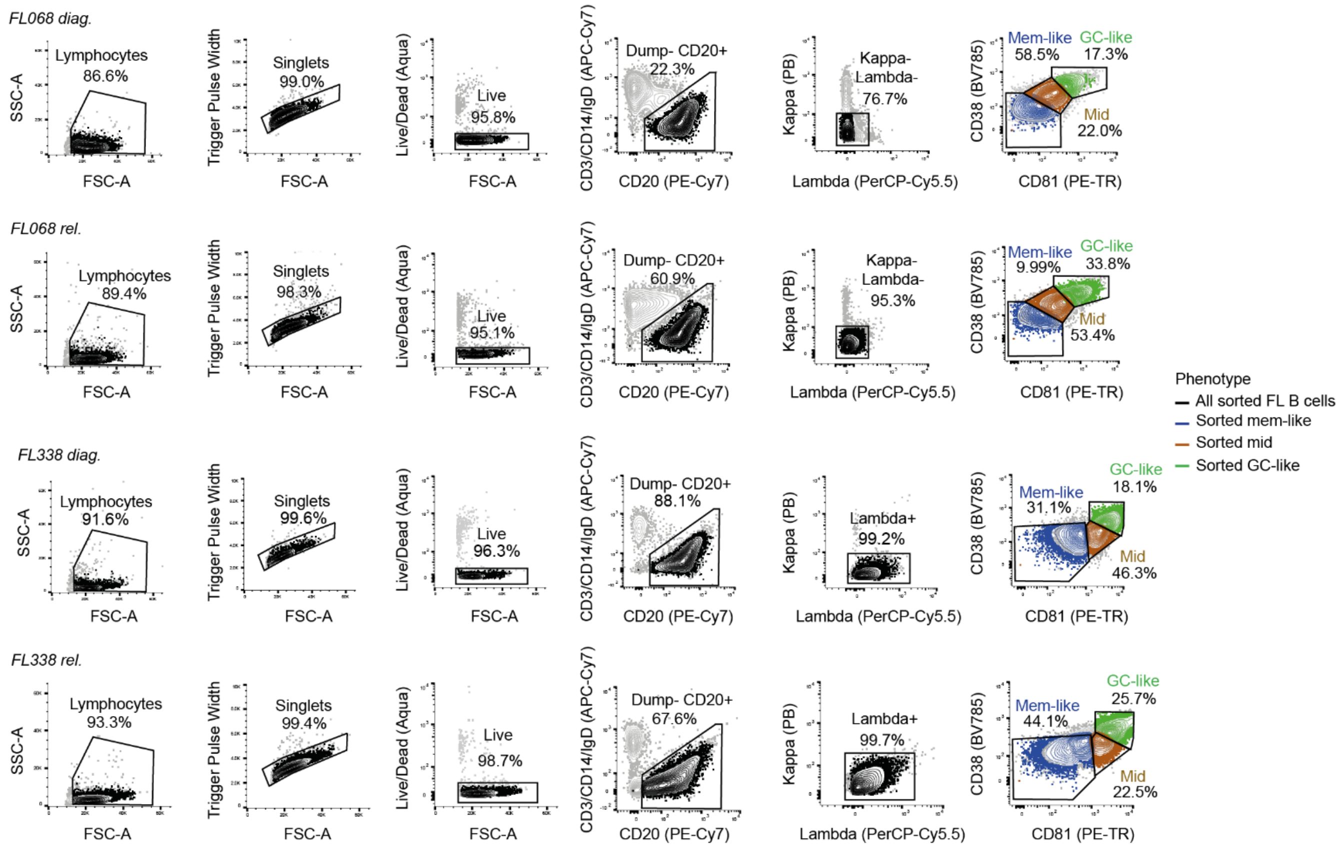


## B

**T Follicular Helper cells sorting strategy for *in vitro* culture**

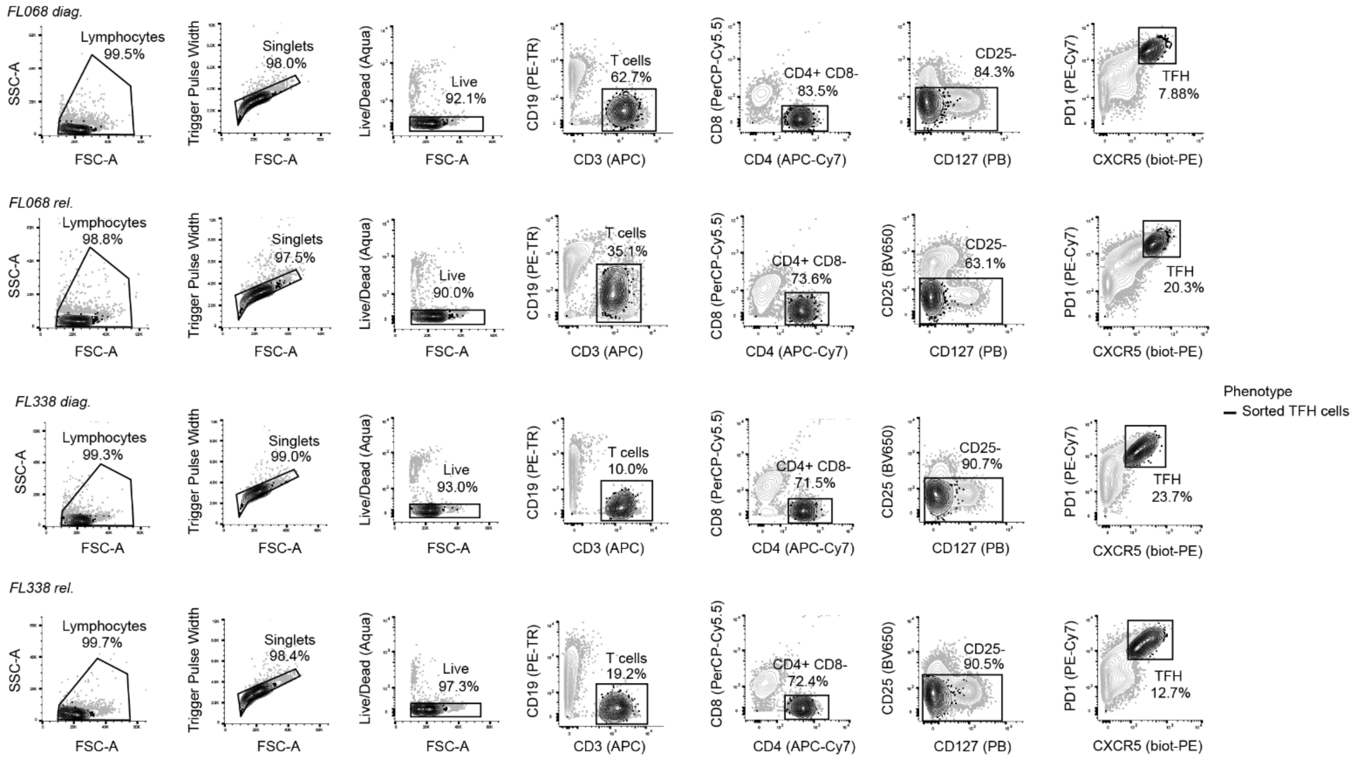

#### Supplementary Figure 7. Cell sorting for *in vitro* culture of GC-like, Mem-like, Mid FL B cells and autologous FL $T_{FH}$ cells

Related to **Fig.6D-E**. **(A)** Flow cytometry gating strategy for bulk cell sorting of GC-like, Mid and Mem-like FL B cells from the indicated samples (FL068 diag., FL068 rel., FL338 diag., FL338 rel.), used for *in vitro* culture experiment. **(B)** Flow cytometry gating strategy for bulk cell sorting of $T_{FH}$ cells from the indicated samples (FL068 diag., FL068 rel., FL338 diag., FL338 rel.). Sorted cells are labeled as colored contours and dots over grey contours and dots (total sample cells), using the color code indicated (Phenotype). Numbers above gates indicate the percentage of cells in the indicated gate relative to its parent gate in the total sample.

#### Supplementary Figure 8

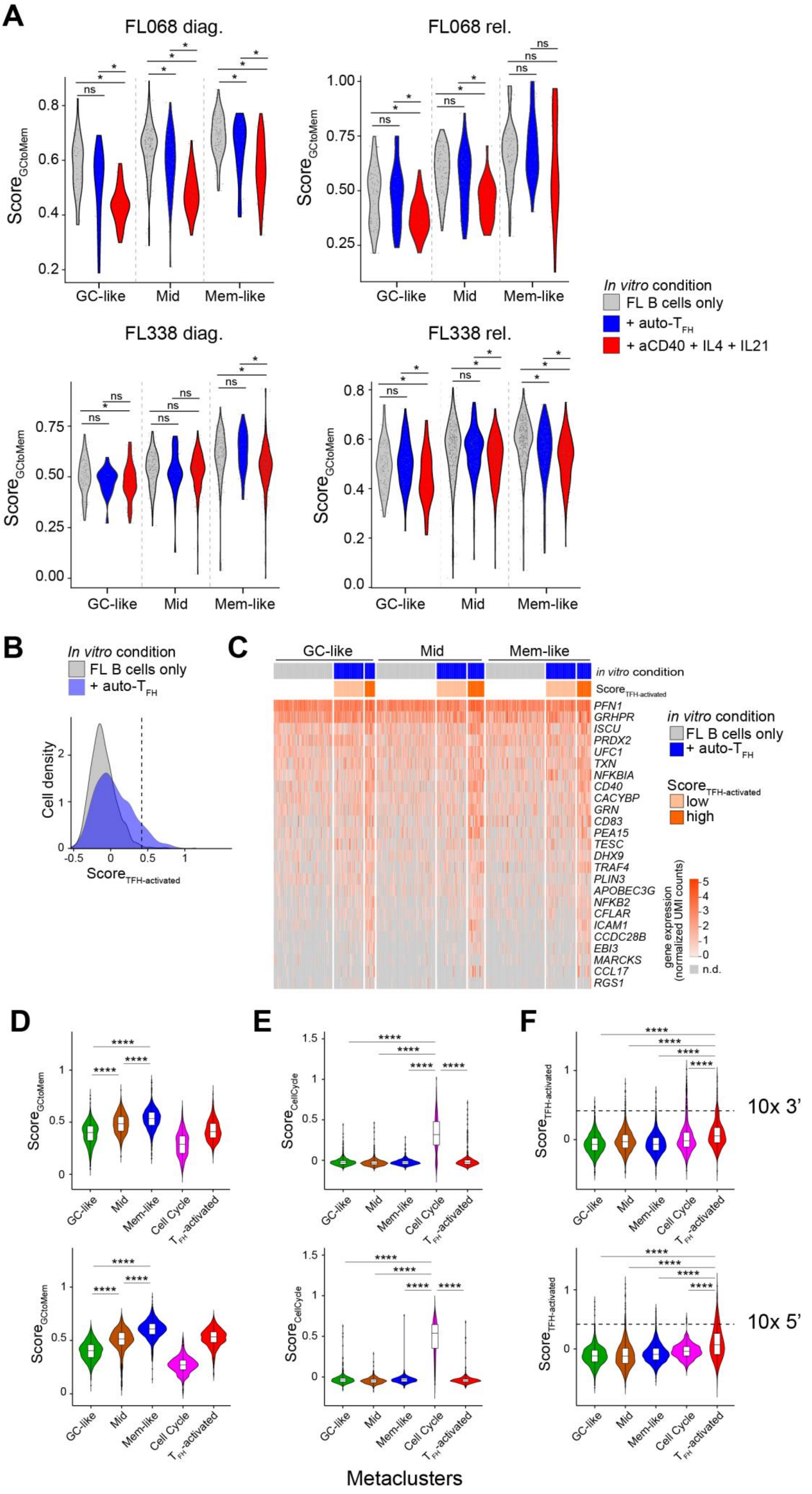

#### Supplementary Figure 8. $T_{FH}$-activated FL B cells

Related to **Fig. 6D-G**. (**A**) Violin plot and dot plot of $Score_{GCtoMem}$ values for sorted GC-like (left), Mid (middle) and Mem-like (right) FL B cells after 48-hour *in vitro* culture in the indicated conditions: FL B cells only (grey), in presence of 1:1 ratio of autologous $T_{FH}$ cells (+ auto-$T_{FH}$, blue), or in presence of $T_{FH}$-derived activation signals (+ aCD40 + IL4 + IL21, red). Each dot represents a single cell. *p<0.05, ns: not significant (Wilcoxon rank sum test with Benjamini & Hochberg adjustment). Each plot corresponds to the indicated sample (FL068 diag., FL068 rel., FL338 diag. and FL338 rel.). **(B)** Density distribution of single-cell $Score_{TFH\text{-}activated}$ values for all FL B cells cultured in absence (FL B cells only, grey) or in presence of 1:1 ratio of autologous $T_{FH}$ cells (+ auto-$T_{FH}$, blue). $Score_{TFH\text{-}activated}$ represents the average expression of genes significantly and recurrently upregulated in FL B cells after culture with autologous $T_{FH}$ cells (genes listed in heatmap **(C)**). Dashed line represents the threshold used to segregate cells with low or high $Score_{TFH\text{-}activated}$ in **(C)**. **(C)** Gene expression heatmap of single GC-like (left), Mid (middle) and Mem-like (right) FL B cells cultured in absence (FL B cells only, first bar, grey) or in presence of 1:1 ratio of autologous $T_{FH}$ cells (+ auto-$T_{FH}$, first bar, blue) for the expression of indicated $T_{FH}$-activated marker genes. Cells from the + auto-TFH condition are further segregated on the basis of low (second bar, beige) or high (second bar, orange) $Score_{TFH\text{-}activated}$ values. **(D-F)** Violin plots of single-cell gene expression scores distributions in FL B cells assigned to the indicated metaclusters from 10x 3' (top) and 10x 5' (bottom) datasets. **(D)** $Score_{GCtoMem}$ values. **(E)** $Score_{CellCycle}$ values. **(F)** $Score_{TFH\text{-}activated}$ values. Dotted line indicates threshold used in **Fig.S8B-C**. **** p < 0.0001 in Wilcoxon pairwise comparisons after Kruskal-Wallis multiple comparisons test.

### Supplementary Figure 9

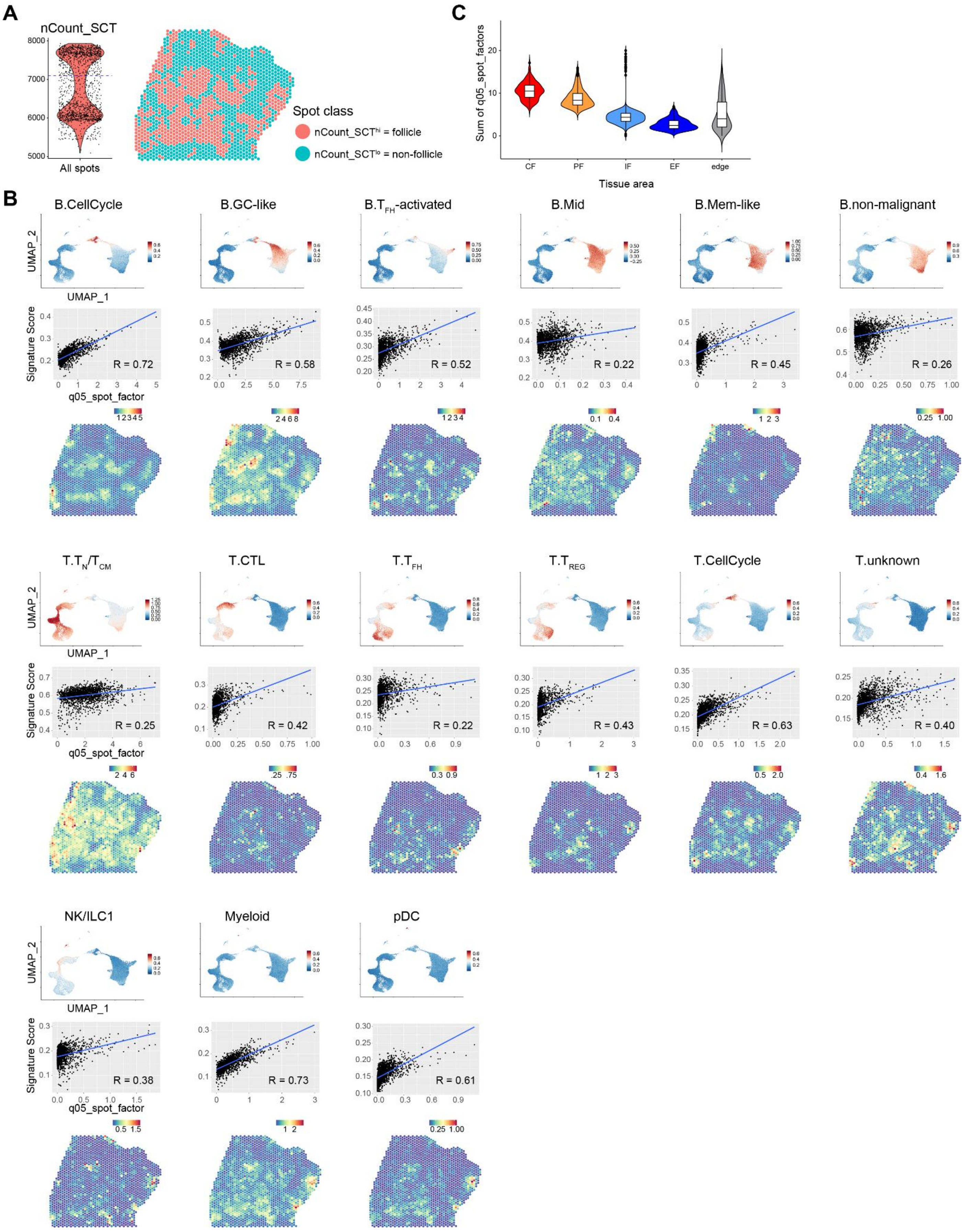

**Supplementary Figure 9. Spatial distribution of malignant B cell subsets in FL lymph node**

Related to **Fig.7**. **(A)** Left: dot plot of normalized gene counts (SCT normalization) in all spots from the spatial transcriptomics data set after gating the region of interest in **Fig.7A**. The dashed blue line indicates the threshold used to classify $nCount\_SCT^{hi}$ follicular spots and $nCount\_SCT^{lo}$ non-follicular spots. Right: spatial transcriptomics spots map, colored by follicular and non-follicular spot class. **(B)** For each cell subset defined in **Fig.7C**, as indicated. (top) UMAP embedding of the integrated scRNA-seq datasets colored by signature score of that cell subset’s marker genes. (middle) Correlation scatter plot of the signature score of that cell subset’s marker genes computed on the spatial transcriptomics spots (y-axis) versus the q05_spot_factors for that subset obtained from deconvolution (x-axis) for spatial transcriptomics spots (blue line: linear regression line, Pearson’s correlation factor as indicated). (bottom) Spatial transcriptomics spatial map colored by q05_spot_factor for that cell subset. **(C)** Violin plot of the sum of q05_spot_factors for all deconvoluted subsets in spatial transcriptomics spots grouped by tissue area defined in **Fig.7B**.

#### Supplementary Figure 10

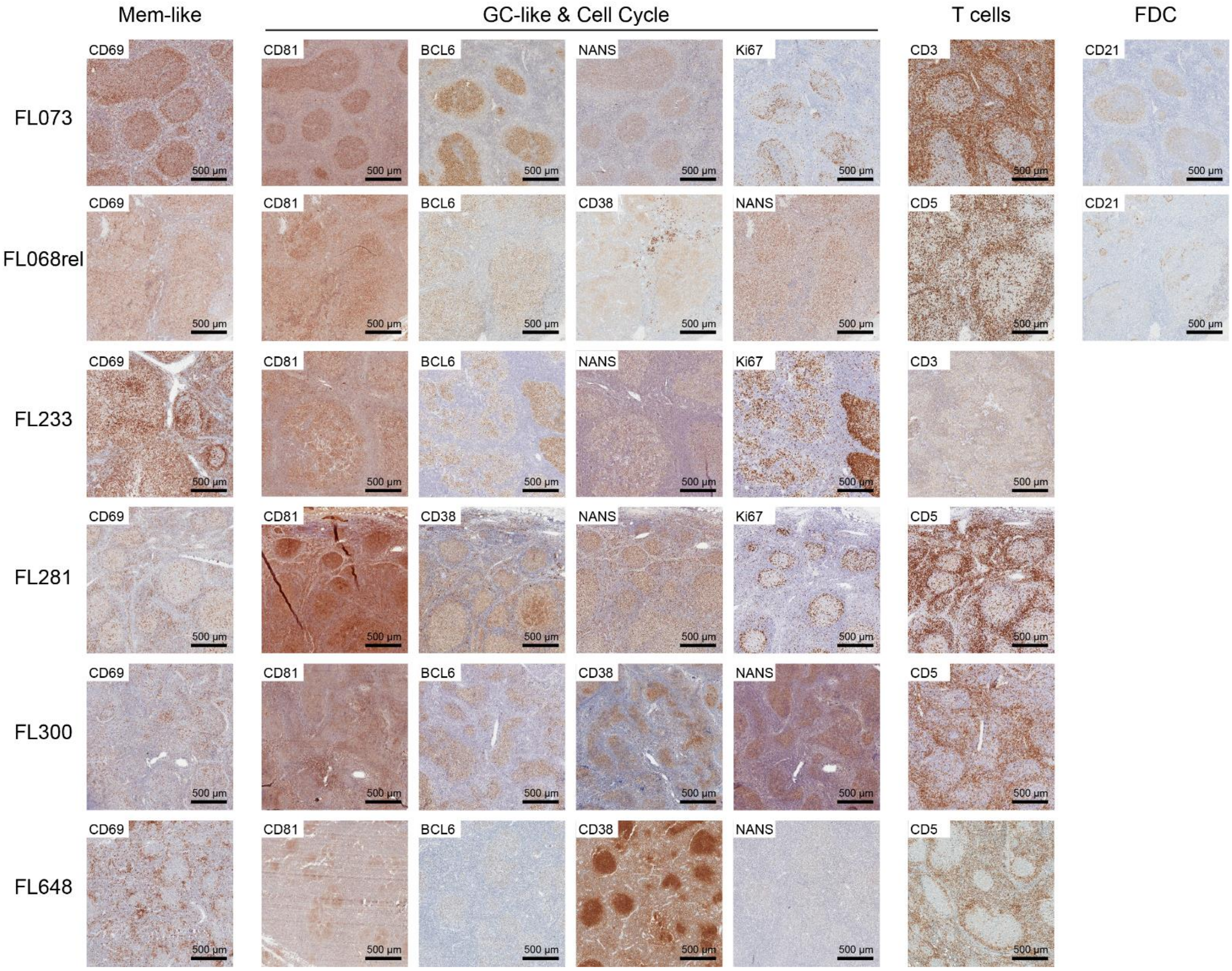


**Supplementary Figure 10. Immunohistochemical analyses of FL lymph nodes**

Related to **Fig.7**. Immunohistochemistry detection of markers for Mem-like (CD69), GC-like and Cell Cycle (CD81, BCL6, NANS, CD38, Ki67), T cells (CD3, CD5) and FDC (CD21) in sections from the indicated FL lymph node biopsy samples. Scale bars = 500µm.
